# Preventing Data Leakage in Neural Decoding

**DOI:** 10.64898/2026.01.26.701583

**Authors:** Robert Wong, Michael H. McCullough, Tianfeng Lu, Shuyu I. Zhu, Geoffrey J. Goodhill

**Affiliations:** Department of Electrical and Systems Engineering, Washington University in St. Louis; Departments of Developmental Biology and Neuroscience, Washington University in St. Louis; Eccles Institute of Neuroscience, John Curtin School of Medical Research, College of Science and Medicine, The Australian National University; School of Computing, College of Systems and Society, The Australian National University

## Abstract

Neural decoding is a widely-used machine learning technique for investigating how behavior, perception and cognition are represented in neural activity. However without careful application data leakage can occur, where information from the test set contaminates the training set, leading to biased estimates of decoding performance and potentially invalidating biological conclusions. Here we show that leakage can be introduced in neural decoding studies by common supervised and unsupervised preprocessing steps, including dimensionality reduction. We reveal that in some cases leakage can paradoxically *decrease* decoding performance relative to unbiased estimates, and we provide theoretical analyses explaining how this occurs. We also show that, for autocorrelated neural time series, randomized k-fold cross-validation can dramatically overstate decoding performance. Based on these findings, we provide detailed recommendations for avoiding data leakage in neural decoding.

## 1 Introduction

Neural decoding is an important tool for understanding how neural activity relates to behavior, cognition, and perception (Horikawa et al., 2013; Anumanchipalli et al., 2019; Glaser et al., 2020; Stringer et al., 2021). It is also widely used for neural engineering applications, including visual decoding and reconstruction from brain activity (Horikawa & Kamitani, 2017; Wen et al., 2017; Chen et al., 2023a,b) and brain-computer interfaces for communication, movement control, and sensory feedback (Wolpaw & McFarland, 2004; Birbaumer & Cohen, 2007; He et al., 2020; Flesher et al., 2021). Recent advances in recording techniques in animals now allow many thousands of neurons to be recorded simultaneously (Ahrens et al., 2013; Jun et al., 2017; Saxena & Cunningham, 2019; Steinmetz et al., 2021; Lin et al., 2022; Urai et al., 2022), resulting in complex and high-dimensional data. Decoding analyses of such data often require sophisticated machine learning methods to translate these measurements into meaningful insights about neural representations and function.

However such analyses risk data leakage, which occurs when information used to train the model also appears in the data used to evaluate performance, even in indirect or inadvertent ways. This can often inflate, and sometimes counterintuitively deflate, the decoding performance of the model and potentially alter the biological interpretation of the results. Data leakage is increasingly recognized as a critical and widespread problem in many areas of science where machine learning techniques are applied, contributing considerably to reproducibility problems (Ambroise & McLachlan, 2002; Simon et al., 2003; Moscovich & Rosset, 2022; Kapoor & Narayanan, 2023). This includes the classic “double-dipping” problem: reusing the same data for feature or region selection and for evaluation, which inflates effect sizes and invalidates inference; selection and evaluation must rely on statistically independent data (Kriegeskorte et al., 2009; Button, 2019). Although train-test separation is now routine in many fields, this split does not prevent leakage if data-dependent preprocessing is applied before the split. For example, Moscovich & Rosset (2022) found that a substantial fraction of papers in a cross-validation (CV) literature survey appeared to apply unsupervised preprocessing before separating training and test data. Recent work has documented leakage in practical fMRI prediction analysis pipelines (Poldrack et al., 2020; Rosenblatt et al., 2024). Yet, despite the growing use of preprocessing procedures for large neural and behavioral datasets where data leakage can easily occur, the broader impact of leakage on neural decoding remains under-recognized.

One route to leakage is data-dependent preprocessing, in which information from held-out samples can shape transformations applied before model fitting. Such preprocessing pipelines can be broadly categorized into two types: supervised and unsupervised. Supervised preprocessing uses label or task information to choose features or fit transformations before decoding, for example selecting neurons based on responsiveness to the decoded variable, selecting features by their correlation with the behavioral outcome, or applying Principal Component Analysis (PCA) to label-defined trial averages. Statistical learning textbooks emphasize that any supervised transformation should be fit exclusively on training data within each CV fold to avoid contaminating test estimates (Hastie et al., 2009; Abu-Mostafa et al., 2012). This is a recurrent pitfall that almost universally leads to optimistic performance estimates if not handled correctly within CV folds. However, a more subtle issue is unsupervised preprocessing, which uses only the neural activity for transformations such as dimensionality reduction or feature scaling. While often considered safe (Hastie et al., 2009), recent theoretical work shows that applying these methods to the entire dataset before partitioning introduces a systematic bias (Moscovich & Rosset, 2022). Crucially, and counter-intuitively, this bias is not universally optimistic; the estimated error can also be pessimistic, i.e. leading to an underestimation of a model’s true decoding performance, depending on a complex interplay between the data, the preprocessing method, and the learning algorithm. A second route to leakage is temporal dependence, where test samples are not statistically independent of training samples (Kapoor & Narayanan, 2023). This issue is especially relevant for neural recordings, where slow internal-state signals and autocorrelated activity can make nearby training and test samples statistically dependent (Harris & Thiele, 2011; Cowley et al., 2020). Such slow structure can also produce spurious associations when analyzed with methods that assume independent samples (Harris, 2021).

Here, we present a systematic analysis of neural decoding pipelines that span supervised and unsupervised preprocessing, modern dimensionality-reduction methods, and static and temporally structured data, benchmarking cross-validated estimates against true generalization. Beyond prior work on feature selection and general-purpose machine-learning pipelines, we focus on preprocessing operations that are central to systems neuroscience, demonstrating that dimensionality reduction can bias decoding performance estimates in either direction. We also derive new analytical results showing how trial-averaged PCA can create separability under random labels, how leaked test samples alter squared-bias and variance, and how train–test covariance biases test error on temporally correlated data.

Together, these results demonstrate that data leakage can occur surprisingly easily in neural decoding. We provide recommendations and open-source code to aid researchers in avoiding data leakage, and also suggest a simple method for detecting the presence of data leakage.

## 2 Results

### 2.1 The role of neural decoding and the challenge of data leakage

Neural decoding treats the relationship between population activity and stimuli or behavior as a supervised prediction task (Fig. 1a): a model is trained on neural activity to predict labels (Fig. 1b) and evaluated on held-out data (Fig. 1c), with performance interpreted as evidence that the population carries information about the target variable (Borst & Theunissen, 1999; Dayan & Abbott, 2005). Performance is typically estimated with CV, which partitions data into train/test (and sometimes validation) sets (Fig. 1d,e) under an assumption of independence between what is used for training and what is used for evaluation. Data leakage violates this assumption when information from the test set influences any stage of model training. A frequent source of leakage is preprocessing performed on the entire dataset before splitting (Fig. 1f). The correct procedure is to split first, then fit all data-dependent preprocessing and model parameters using only the training data in each fold, and finally apply the learned transformations to the test data (Fig. 1g). While the potential for data leakage in supervised preprocessing methods (which use labels) is relatively clear, surprisingly, leakage can also occur from unsupervised methods. We now discuss both in more detail. Throughout, we use “sample” to mean one observation supplied to the preprocessor or decoder, and “trial” to mean one experimentally defined episode, such as a stimulus presentation, choice, or behavioral run. In stimulus-evoked analyses a trial often contributes a single sample, whereas in continuous behavior or navigation tasks a single trial may contain many time-resolved samples.

**Figure 1:**
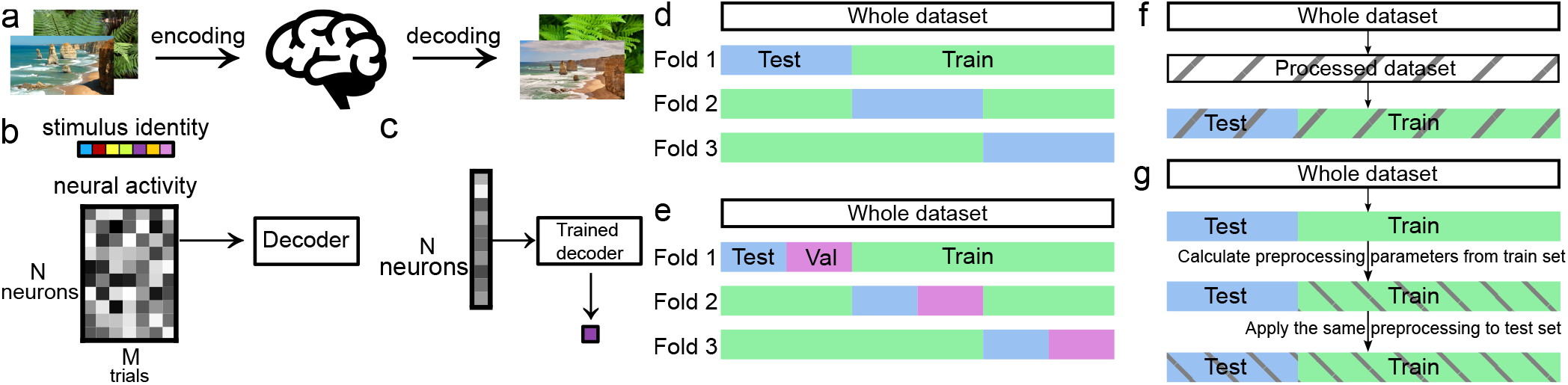
The neural decoding framework and the importance of a non-leaky cross-validation (CV) pipeline. **(a)** Schematic of neural encoding and decoding. Encoding is the process by which a stimulus generates a neural response. Decoding is the inverse problem of inferring the stimulus from the observed neural response. **(b)** The training phase of neural decoding. A decoder (machine learning model) learns a mapping from a training dataset, consisting of neural activity patterns (N neurons over M trials), to their corresponding stimulus or behavioral labels. **(c)** The evaluation phase. The trained decoder is applied to a new, held-out test sample of neural activity to predict its corresponding label. The accuracy of this prediction is used to quantify decoder performance. **(d)** Standard CV data partitioning. A dataset is split into training (green) and test (blue) sets over multiple folds. **(e)** An optional validation set (mauve) can be used for hyperparameter tuning. **(f)** An incorrect, “leaky” data pipeline. Preprocessing is applied to the whole dataset (grey hatching across the full bar) and the resulting processed data are then split into training and test sets (green and blue segments with the same hatching), so the test data influence the learned preprocessing and contaminate the training set, leading to biased performance estimates. **(g)** A corrected non-leaky data pipeline. The dataset is first partitioned into training and test folds. Preprocessing parameters are then learned only from the training data (green segment with hatching) and the same transformation is applied to the held-out test data (blue segment with matching hatching), ensuring that test data do not affect how preprocessing is learned.

### 2.2 Data leakage via supervised preprocessing

#### 2.2.1 Supervised feature selection

Feature selection aims to retain neurons informative about the variable of interest while discarding noisy or uninformative neurons. Common supervised approaches include fitting tuning curves, computing neuron-label correlations, regression coefficients or ANOVA F-statistics; additional methods include L1 (lasso) regularization, tree-based models, and sequential selection (Ferri et al., 1994; Tibshirani, 1996; Lotte et al., 2018). Although valuable in the settings of high-dimensional neural data, applying any label-dependent selection step to the full dataset before CV can induce data leakage (Poldrack et al., 2020; Shim et al., 2021; Rosenblatt et al., 2024), and in at least one case, contributed to a retraction (Verstynen & Kording, 2023).

Here, “feature selection” refers to the post hoc rule used to decide which recorded neurons enter the decoder (not e.g. to whether a neuron was physically present on the electrode or in the imaging plane). Leakage occurs when the statistic used for this decision is estimated from the same samples that will later be treated as held out. We first illustrate this with a simulation. Neural responses to two stimuli were generated from multivariate Gaussian distributions with matched covariance and slightly different means (two-neuron example illustrated in Fig. 2a), each neuron had a Gaussian tuning curve with the same tuning width but a different tuned stimulus. The theoretical optimal classification accuracy in this case is 69% (Methods). We increased the number of recorded neurons while keeping the total number of stimulus presentations fixed, mimicking trial-limited experiments. To remove potentially noisy neurons and reduce computation time, feature selection was employed to select high-quality neurons (Fig. 2b). ANOVA F-statistics were computed separately for each neuron, then used to rank the population and select the top 100 neurons before 5-fold CV. When this ranking was performed on the entire dataset, the selected set could change when held-out trials were included. Decoding accuracy then increased with dimensionality but also exceeded the theoretical optimum (Fig. 2c). The same qualitative inflation occurred without the Gaussian tuning curve assumption (Fig. S1a). This inflation arose because neurons were chosen using information from all samples, including those later assigned to test folds. Thus, selection was partially driven by noise fluctuations specific to the test data, producing an overly optimistic estimate. Consistent with this, the fold-level decoding accuracy covaried with the amount by which the held-out trials inflated the F-statistic of the selected neurons, and inflation vanished when the ranking was restricted to the training trials (Fig. S2). An additional validation set with nested CV can provide a more accurate estimate of model accuracy when a model requires hyperparameter tuning (Stone, 1974; Cawley & Talbot, 2010). However, this would not rectify the above issue because the contamination occurs before any data partitioning. Performing feature selection strictly within each training fold, and then applying the chosen neurons to the held-out fold, kept decoding accuracy below the theoretical ceiling (Fig. 2d, Fig. S1b).

**Figure 2:**
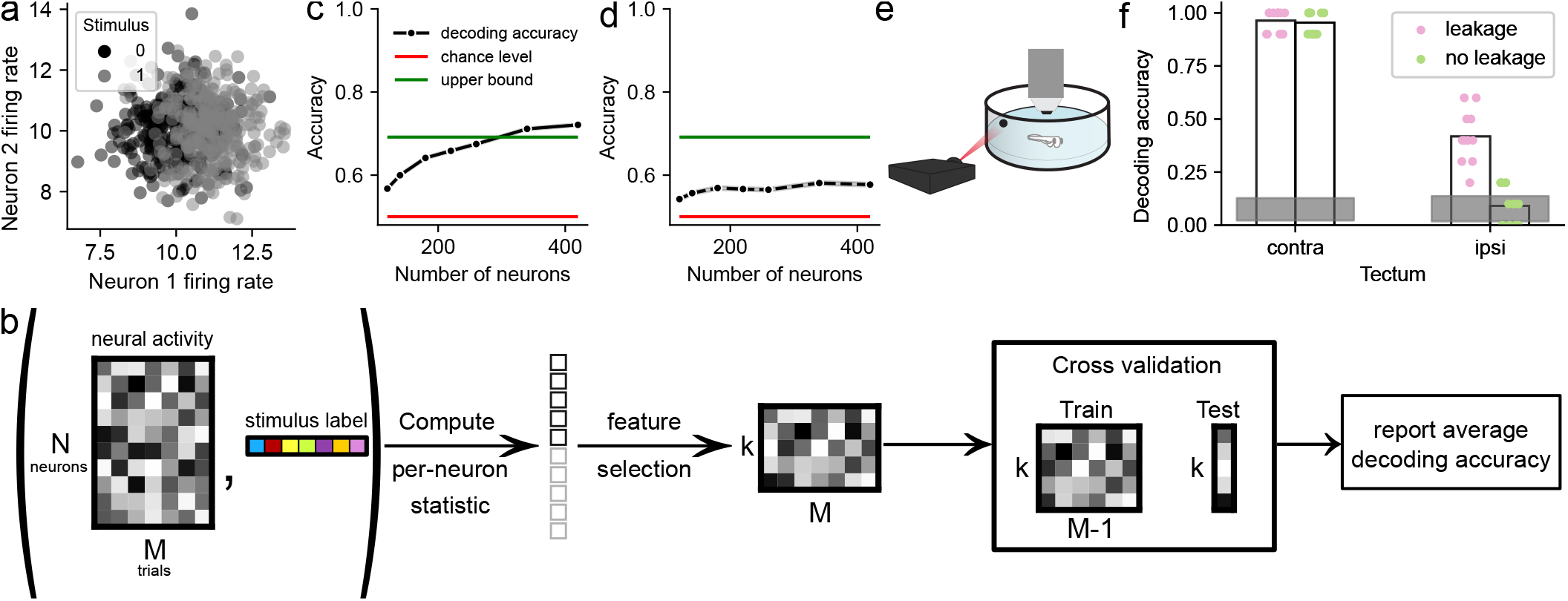
Data leakage through feature selection. **(a)**: Neural activity simulated from a multivariate Gaussian distribution with different class means; example activities in a two-neuron case. Each dot represents one simulated trial, colored by stimulus class. **(b)**: Illustration of the feature-selection method applied to the neural activity matrix and the stimulus identity vector. Neurons were ranked based on the statistical relationship between each neuron’s response to the stimuli and corresponding stimulus identities, and the top k most-informative neurons were kept (the first 5 neurons were kept in the schematic). This reduced dataset was then used for further analysis. A Leave-One-Out CV scheme is illustrated here, where the dataset was divided into a training set (M-1 trials) for training the decoder and a test set (1 trial) for evaluation. The average decoding accuracy in the evaluation phase across all CV folds is reported as the overall accuracy. **(c)**: In a pipeline with data leakage, feature selection leads to inflated performance when used as a preprocessing step on the whole dataset. The horizontal axis shows the number of recorded candidate neurons before feature selection; at each value, the top 100 neurons were selected for decoding. The 95% confidence interval of the mean from 100 independent simulations are too small to be visible. Green line is the theoretical upper bound of decoding accuracy for a linear decoder. **(d)**: In a pipeline without data leakage, feature selection was calculated from training data only, and the same selection was then applied to the test set. Accuracy is lower than in (c) and remains within theoretically-achievable bounds. **(e)**: Schematic of a larval zebrafish embedded in agarose for an imaging experiment. The fish was shown stationary visual stimuli at angles from 15°to 165°in increments of 15°. **(f)**: In a pipeline with data leakage, both sides of the tectum showed good decoding. However after ensuring no data leakage, decoding accuracy in the ipsilateral tectum fell to chance level, indicating that no relevant stimulus information is being encoded in this region. Each dot is the averaged accuracy from 10 CV loops. Shaded region: 95% confidence region from shuffle control.

We next examined a real-world example from the visual system. Two-photon calcium imaging of zebrafish larvae was used to record tectal activity at single-cell resolution in response to spot stimuli presented at discrete locations to the right side of the visual field (Fig. 2e; see Methods). We asked how well spot position could be decoded from neural activity in both contralateral (left) and ipsilateral (right) tecta. In zebrafish the retinotectal projection is completely crossed, and so the ipsilateral tectum receives no direct retinal input. Again ANOVA F-statistics were computed on the entire dataset to select the top 100 neurons, a linear decoder was then trained on their neural activity, and the accuracy from CV reported. Unsurprisingly decoding performance in the contralateral tectum was very good (Fig. 2f). However decoding performance in the ipsilateral tectum was also good (Fig. 2f). Recent work has identified a population of intertectal neurons in zebrafish that transmit information between hemispheres (Gebhardt et al., 2019), providing a possible substrate for the transfer of spatial information to the ipsilateral tectum. However alas the decoding result is wrong, for the same reasons as in our simulated example: the test set’s information influenced the selection of features, leading to a biased training process where the model ‘learned’ from the data it was supposed to predict. When the decoding analysis was repeated without data leakage – by deriving the features to be selected only on the training set – decoding accuracy in the ipsilateral tectum fell to chance level (Fig. 2f), indicating that no spatial information was transferred. Analogous issues for label-free criteria, such as variance- or firing-rate-based selection, are treated separately below as examples of unsupervised feature selection.

#### 2.2.2 Supervised dimensionality reduction

Dimensionality reduction is often used before decoding because neural recordings are commonly high-dimensional relative to the number of trials. Restricting the decoder to a lower-dimensional representation can denoise shared population activity, reduce estimator variance, and act as a form of regularization (Cunningham & Yu, 2014; Mignacco et al., 2020; Green & Romanov, 2025). Empirically, several neural population studies have used supervised dimensionality reduction before decoding and reported improved predictive performance compared to using the raw population activity in its original dimensionality (e.g. Rumyantsev et al., 2020; Ebrahimi et al., 2022). Supervised dimensionality reduction learns label-informed projections of population activity (Perich et al., 2025). When the projection is fit on the entire dataset before CV, test-set structure shapes the latent space, making conditions appear more separable and distorting reported decoding error estimates.

To show that globally-fit supervised dimensionality reduction inflates empirical performance and that fold-wise fitting removes this underestimation of error, we first simulated neural activity with independently generated neuronal responses and randomly assigned stimulus labels. Trial-averaged PCA (a supervised step because averaging uses labels) was used to obtain low-dimensional embeddings. Here, trial averaging first converts the trial-by-neuron activity into a condition-by-neuron matrix of class means, where each row is the mean response across all trials of one stimulus condition; PCA is then computed on this matrix of condition means, yielding axes of maximal variance across the estimated class means. Individual trials are subsequently projected onto these PC axes for visualization and decoding, so each point in the PC embedding corresponds to one trial (Fig. 3a,d). In this respect, trial-averaged PCA resembles supervised dimensionality-reduction methods that seek low-dimensional axes separating label-defined groups; under isotropic within-condition covariance and balanced conditions, it is closely related to linear discriminant analysis.

**Figure 3:**
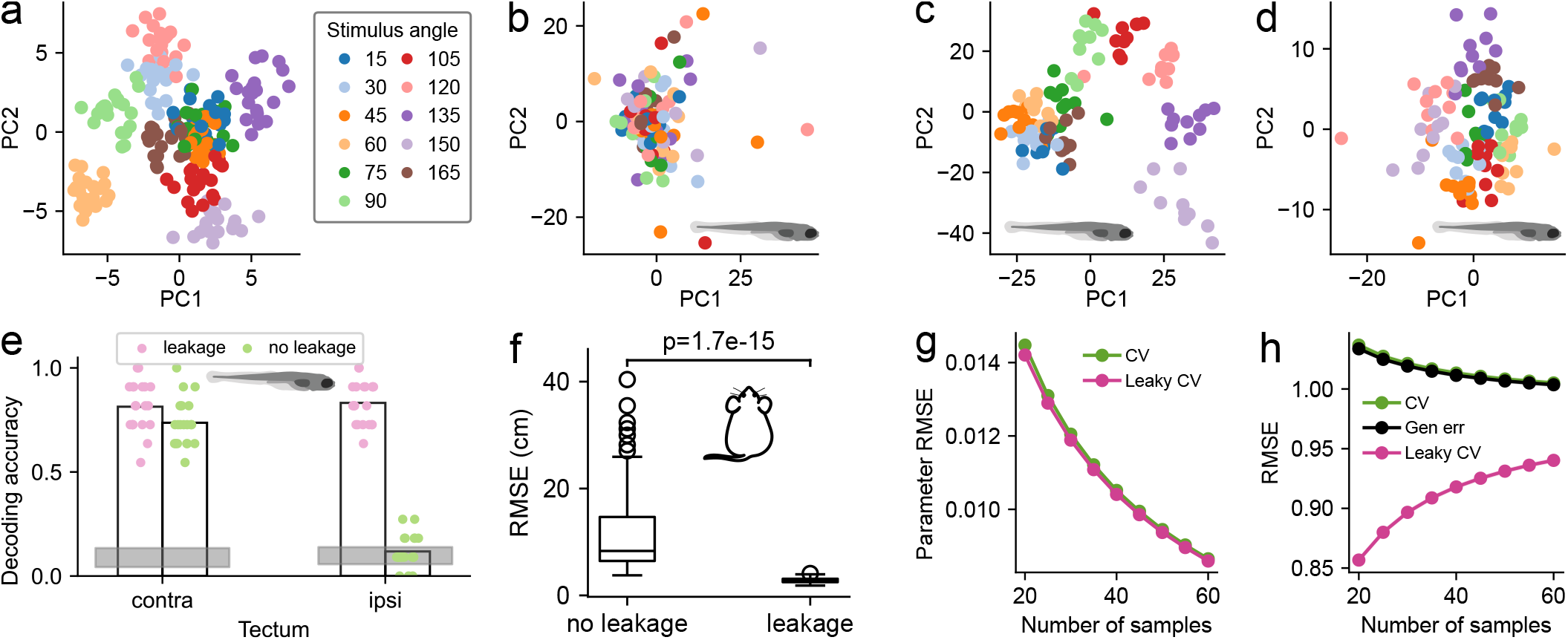
Data leakage through supervised dimensionality reduction. **(a)**: Simulated random neural activity, PC space calculated from trial-averaged data and all trials were projected to this space (see Methods). Each dot corresponds to one trial. Structure is apparent even though there is actually no structure in the data. **(b-c)**: 2D PC embedding of zebrafish tectal activity. (b): ipsilateral tectum, (c): contralateral tectum. The zebrafish (Petrucco, 2020) and rodent icons indicate the animals from which these data were obtained. **(d)**: PC embedding using trial-averaged PCA of the same ipsilateral activity as in (b), showing apparent clusters of neural activity responding to the same stimulus. **(e)**: Decoding in a pipeline with data leakage suggests high signal fidelity encoded in both ipsilateral and contralateral tectum. However decoding accuracy falls back to chance level for the ipsilateral tectum in a pipeline without data leakage. Each dot is the averaged accuracy from one CV loop. Shaded region: 95% confidence region from shuffle control in a pipeline without data leakage. **(f)**: Supervised CEBRA (Schneider et al., 2023) was used to reduce the dimensionality from the recorded neural activity and to decode a rat’s location in a linear track. Decoding accuracy is inflated in a pipeline with data leakage compared to a pipeline without leakage. p-value calculated from two-sided Wilcoxon signed-rank test, n=84. **(g**,**h)**: With simulated data and ground truth parameters, partial least squares (PLS) with one component was used for supervised dimensionality reduction. **(g)**: Although the leaky CV includes one more leaked sample point in the dimensionality reduction step, the parameter RMSE is similar between the leaky and non-leaky cases. SEMs across 10,000 simulation runs are too small to be visible. **(h)**: The CV RMSE from the leaky pipeline underestimates the true generalization error of the resulting regression model. Non-leaky CV matches the true generalization error, despite fitting dimensionality reduction with one fewer sample per fold.

Despite no true signal, when the number of neurons greatly exceeded the number of samples, global trial-averaged PCA manufactured spurious clusters and apparent structure (Fig. 3a). The degree of spurious clustering, quantified by the multiclass Fisher discriminant ratio, was governed by the ratio between the number of neurons and number of trials per condition (Fig. S4), and could be predicted analytically (Section S2). Applying this to the tectum data used in Fig. 2, standard PCA showed no stimulus information in ipsilateral tectum (Fig. 3b) but clear separation in contralateral tectum (Fig. 3c). Computing trial-averaged PCs globally and then training a linear decoder on these embeddings produced visible clustering (Fig. 3d, Fig. S3) and unexpectedly high accuracy (Fig. 3e) for the ipsilateral side—an instance of leakage because all trials contributed to the PCs. Refitting trial-averaged PCA within each training fold and projecting held-out trials restored ipsilateral decoding to chance (Fig. 3e).

We then analyzed CA1 data related to position (Grosmark & Buzsáki, 2016), embedding neural activity with supervised CEBRA (Schneider et al., 2023) and decoding with k-nearest neighbors. Fitting CEBRA globally before CV yielded low test root mean squared error (RMSE), whereas fold-wise fitting increased RMSE to a more realistic level (Fig. 3f).

Two possible reasons could explain the optimistic RMSE estimates observed when supervised projections are fit globally. First, leaked held-out points could improve the fitted model by helping it estimate a more accurate projection. Second, the model itself might not improve, but the reported CV error could become optimistic because the evaluation points had already influenced the projection used to predict them. To distinguish between these possibilities, we used a simulated linear regression problem with known ground-truth coefficients ***β*** and one-component Partial Least Squares (PLS) as the supervised dimensionality-reduction step. For centered data, this PLS step learned a one-dimensional projection from the covariance between features and labels; a linear regressor was then fit on the projected training data. For simplicity, in the analysis below we considered the case in which one point was held out at a time in each CV loop. Across training sample sizes, we compared the following three diagnostic quantities: (i) leaky CV, where the PLS projection was fit using the training data plus the held-out test point, but the linear regressor was fit only on the projected training data; (ii) non-leaky CV, where both PLS and the regressor were fit using training data only and then evaluated on held-out points; and (iii) the true generalization error of the model from (i), evaluated on independent new samples that were not used to fit either the PLS projection or the regressor. Because the true ***β*** was known in this simulation, we also measured parameter RMSE: the error between the estimated regression coefficients, mapped back to the original feature space, and ***β***. This ground-truth comparison directly measured coefficient recovery, whereas CV RMSE measured prediction error on samples designated as held out by the CV split.

These results support the second explanation above, rather than genuine improvement in the estimator. The leaky and non-leaky pipelines produced nearly identical parameter RMSE (Fig. 3g), indicating that the leaked point did not materially improve recovery of the known generative coefficients. However, the leaky CV RMSE underestimated the true generalization error of that same leaky model, whereas non-leaky CV aligned with the generalization-error curve (Fig. 3h). Thus, the apparent improvement in leaky CV error arose from violated independence between preprocessing and evaluation, not from a better estimator. All three curves converged to a common asymptote as the sample size grows: the population prediction error of the 1-component PLS–regression model. At finite sample sizes they approached this limit from opposite sides. The true generalization RMSE and non-leaky CV RMSE sit above the asymptote because finite-sample estimation variance in the fitted regression coefficient inflates prediction error. The leaky CV RMSE sits below the asymptote because the held-out point’s contribution to the fitted PLS direction biases the prediction toward its own label. Both effects shrink with increasing sample size, but at the sample sizes typical of decoding experiments the resulting gap is large enough to materially distort error estimates (Fig. 3h). Thus, supervised dimensionality reduction must be fit fold-wise; global fitting corrupts validation metrics and misguides hyperparameter tuning, and can therefore increase error on future held-out data.

### 2.3 Data leakage via unsupervised preprocessing

A critical, yet often overlooked, form of data leakage can arise during unsupervised preprocessing from the features (neural activity) themselves. Standard methods in data analysis pipelines, such as dimensionality reduction (e.g., PCA), feature scaling, or variance-based feature selection, can introduce error-estimation bias if they are applied to the entire dataset before partitioning for CV (here and elsewhere unless otherwise noted, error-estimation bias refers to the mismatch between a cross-validated error estimate and the true generalization error of the same model, distinct from the squared-bias term in the bias–variance decomposition below.) The application of unsupervised methods across the full dataset is a more insidious pitfall, partly because of contradictory suggestions in leading statistics textbooks (Hastie et al., 2009, p. 246 vs. Abu-Mostafa et al., 2012, p. 174). Recent theoretical work provides a rigorous foundation for understanding this problem. In particular, Moscovich & Rosset (2022) demonstrated analytically that applying data-dependent unsupervised transformations globally can make the cross-validated error estimate differ systematically from the model’s true generalization error. Crucially, they showed that this mismatch is not universally optimistic. Its sign and magnitude depend on an intricate interplay between the data distribution, the specific preprocessing method, the subsequent modeling algorithm, and the CV parameters. We next examined this bidirectional (optimistic or pessimistic) deviation from true generalization error across common unsupervised steps in neural decoding: feature selection, z-scoring, PCA, and modern unsupervised dimensionality reduction algorithms. A bias–variance decomposition of Principal Component Regression (PCR) serves as a case study to explain when and why the sign of the leakage-induced mismatch reverses.

#### 2.3.1 Unsupervised feature selection

To investigate the practical impact of this phenomenon in a common neural decoding context, we simulated neural responses to a circular stimulus variable (e.g., orientation or direction of motion) using the Gamma-Poisson generative model of Pillow & Scott (2012) (Fig. 4a). In the main simulation, each neuron had a Gaussian tuning curve over stimulus angle, so stimulus-dependent modulation was defined as part of the response-generation model. We first confirmed that without any feature selection, a simple linear decoder could be trained to effectively predict the stimulus angle from the population activity, establishing that the decoding problem is well-posed (Fig. 4b). We then implemented a natural preprocessing workflow for high-dimensional neural recordings: we performed unsupervised feature selection by selecting the subset of neurons with the highest spike count variance across the entire dataset before CV. This criterion is intuitive, as neurons with higher variance are generally those whose firing rates are more strongly modulated by the different stimuli, making them more informative for decoding. However, when this ranking is computed once on the full dataset, held-out trials have already influenced which neurons enter the decoder. The procedure is therefore leaky even though it does not use stimulus labels.

**Figure 4:**
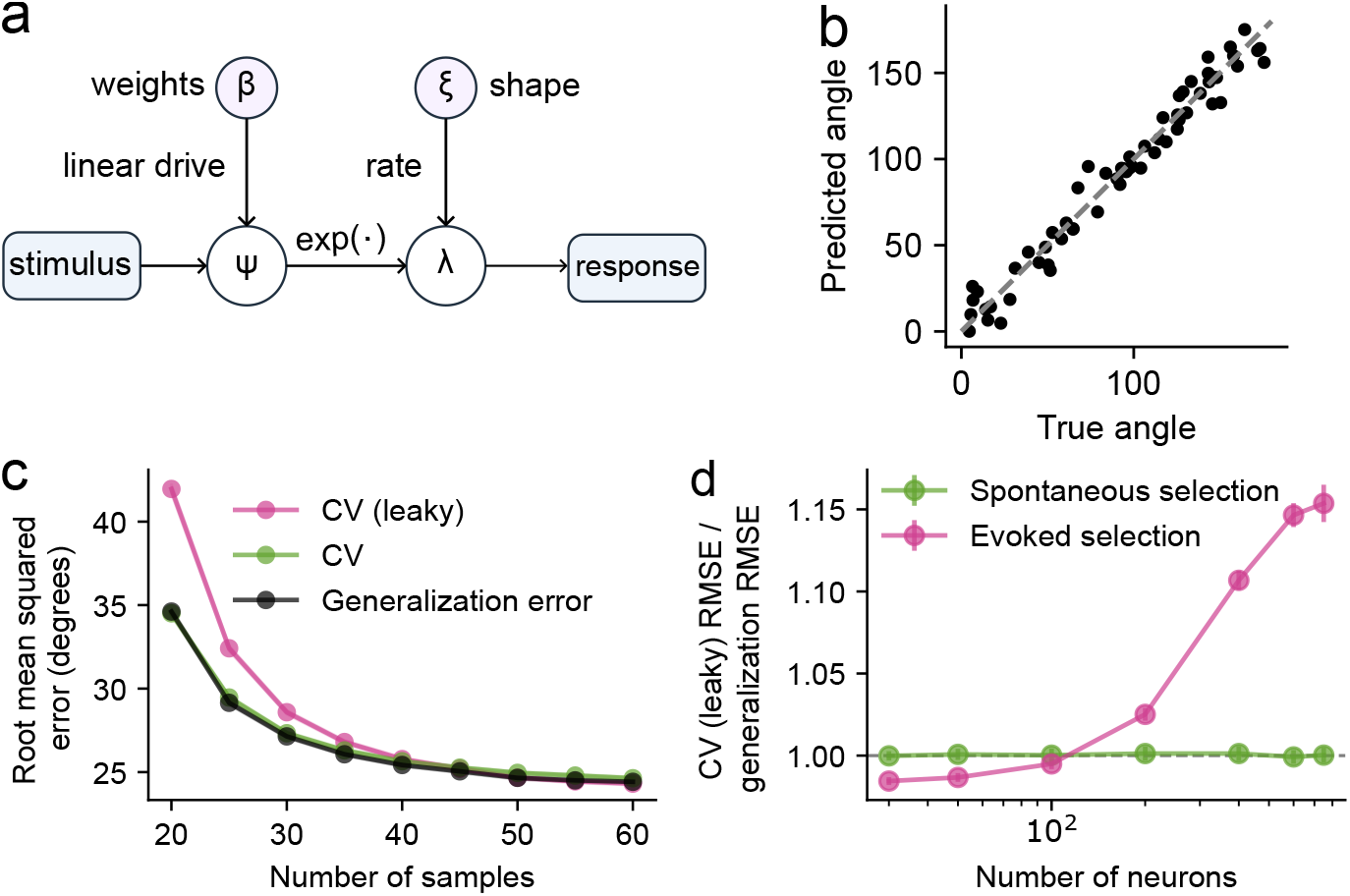
Unsupervised feature selection shows biased results compared to true generalization error. **(a)**: Schematic of the generative model for simulating Poisson-like spike responses of stimuli, adapted from Pillow & Scott (2012). **(b)**: A simple linear decoder achieved good decoding results on held-out data. **(c)**: However when unsupervised feature selection was applied to select a subset of neurons (ranked by variance) for decoding, the CV estimate of decoding error became biased with respect to the true generalization error. In this case the result from data leakage harmed decoding performance (lower root mean squared error means better decoding), thus yielding a more pessimistic estimation. **(d)** In a trial-limited neuron-count sweep using 100 samples and 5-fold CV, the top 10% of neurons were selected by spike-count variance. Selection based on evoked activity, which included data from the held-out folds, increased the leaky-CV/generalization RMSE ratio as the recorded population grew. Selection based on stimulus-independent spontaneous activity served as a non-leaky control and remained close to one. Lines show means across 10,000 simulation runs; uncertainty bands or error bars denote *±*3 SEM when visible.

The leaky model’s estimated error from the CV procedure consistently deviated from its true generalization error (Fig. 4c). Intuitively, this mismatch reflected a bias–variance trade-off: scoring neurons on the full dataset partially fit the held-out trials’ fluctuations, which lowered apparent squared-bias but inflated prediction variance, with the net sign set by whichever effect dominated. The same phenomenon is considered analytically in Section S4. In contrast, the non-leaky procedure recomputed the variance ranking and fit the decoder using only the training samples in each CV fold. Applying each fold’s selected neurons to its held-out samples produced CV errors that closely tracked true generalization error. This effect was not dependent on the particular unsupervised score or on the parametric form of the tuning curves: mean firing-rate selection and a semi-parametric Gamma-Poisson model without explicit Gaussian tuning assumptions produced the same qualitative separation between leaky CV and true generalization error (Fig. S6a–c). Notably, across these scenarios, data leakage could result in pessimistic or optimistic deviations from true generalization error, where the estimated decoding error from leaky CV could be either higher (worse) or lower (better) than the model’s true performance on new data.

We next asked when this unsupervised-selection leakage becomes large enough to matter in practice. We compared two variance-based ranking rules by performing a Gamma-Poisson neuron-count sweep with 100 samples, 5-fold CV, Gaussian tuning curves, and selection of the top 10% of neurons. “Evoked” selection ranked neurons using the stimulus-evoked samples that entered the decoding analysis, whereas “spontaneous” selection ranked neurons using stimulus-independent activity outside the evoked decoding samples. With evoked selection, the leaky-CV/generalization error ratio remained close to one for small populations but increased as the total recorded population approached hundreds of neurons (Fig. 4d). In contrast, spontaneous selection kept the error ratio close to one (Fig. 4d). The same qualitative contrast between evoked and spontaneous selection was observed when neurons were ranked by mean firing rate rather than variance (Fig. S6d). The sampling noise in each neuron’s mean or variance estimate is mainly determined by the number of trials, not by the number of recorded neurons. The population-size dependence arises at the ranking step: with more candidate neurons, random fluctuations from the held-out folds have more chances to change which neurons are selected. Thus, the leaky and training-only selected feature sets are expected to be similar in low-dimensional, trial-rich settings, but can diverge in trial-limited, high-dimensional recordings. A simplified Gaussian-feature calculation with an ordinary least squares decoder gives the same qualitative prediction: the ratio is bounded, but increases as the number of selected neurons approaches the number of training samples per fold. For the 100 samples, selecting the top 10% neurons from 750 neurons gives an expected error ratio of approximately 1.12 in this Gaussian benchmark (Section S3). Thus, criteria specified independently of the evoked decoding samples need not be nested within CV, whereas scores computed from those samples should be recomputed within each training fold.

#### 2.3.2 Unsupervised dimensionality reduction

Another common and powerful unsupervised preprocessing step in neural decoding is dimensionality reduction. Unsupervised dimensionality reduction algorithms identify patterns and structures in data without the need for labeled outcomes or predictions (Kriegeskorte & Wei, 2021; Langdon et al., 2023). They can uncover hidden structures and correlations within complex, high-dimensional datasets and are particularly important when dealing with spontaneous neural activity or unlabelled naturalistic behavior, facilitating the discovery of intrinsic neural representations and functional connectivity without prior knowledge.

PCA is a cornerstone of many neural data analysis pipelines, providing a linear mapping from high-dimensional activity to lower-dimensional components (Fig. 5a). We first investigated how data leakage affects PCR, in which PCA precedes linear regression. In a leaky CV pipeline, PCA was fit using the training samples plus the held-out sample in each CV loop, whereas the regression model was fit only on the projected training samples. In the non-leaky CV pipeline, both PCA and regression were fit using only the training samples. The resulting regression parameters had comparable quality between the two pipelines (Fig. 5b), and the sample mean squared error (MSE) from the non-leaky CV closely matched the true generalization error of the leaky pipeline (Fig. 5c). In contrast, the sample MSE obtained from the leaky CV procedure was, surprisingly, a pessimistic estimate of its own generalization error (Fig. 5c,d). To understand this counterintuitive behavior, we decomposed the explainable MSE (excluding the irreducible label noise) into squared-bias and variance terms (Fig. 5e; see Methods and Sections S4.3 and S4.4). When PCA was fit with leakage, the squared-bias term in the MSE decomposition was systematically underestimated (optimistic) while the variance term was overestimated (pessimistic). Because the leaky PCA subspace was more aligned with test data, it reduced apparent squared-bias but inflated variance. Thus, the overall explainable MSE, which combines opposing squared-bias and noise-scaled variance contributions, can be either optimistic or pessimistic depending on the regime (Fig. 5f–h). This dependence also varied with sample count: in low-trial regimes the CV MSE could overestimate true generalization error, an effect that diminished with more trials (Fig. 5e).

**Figure 5:**
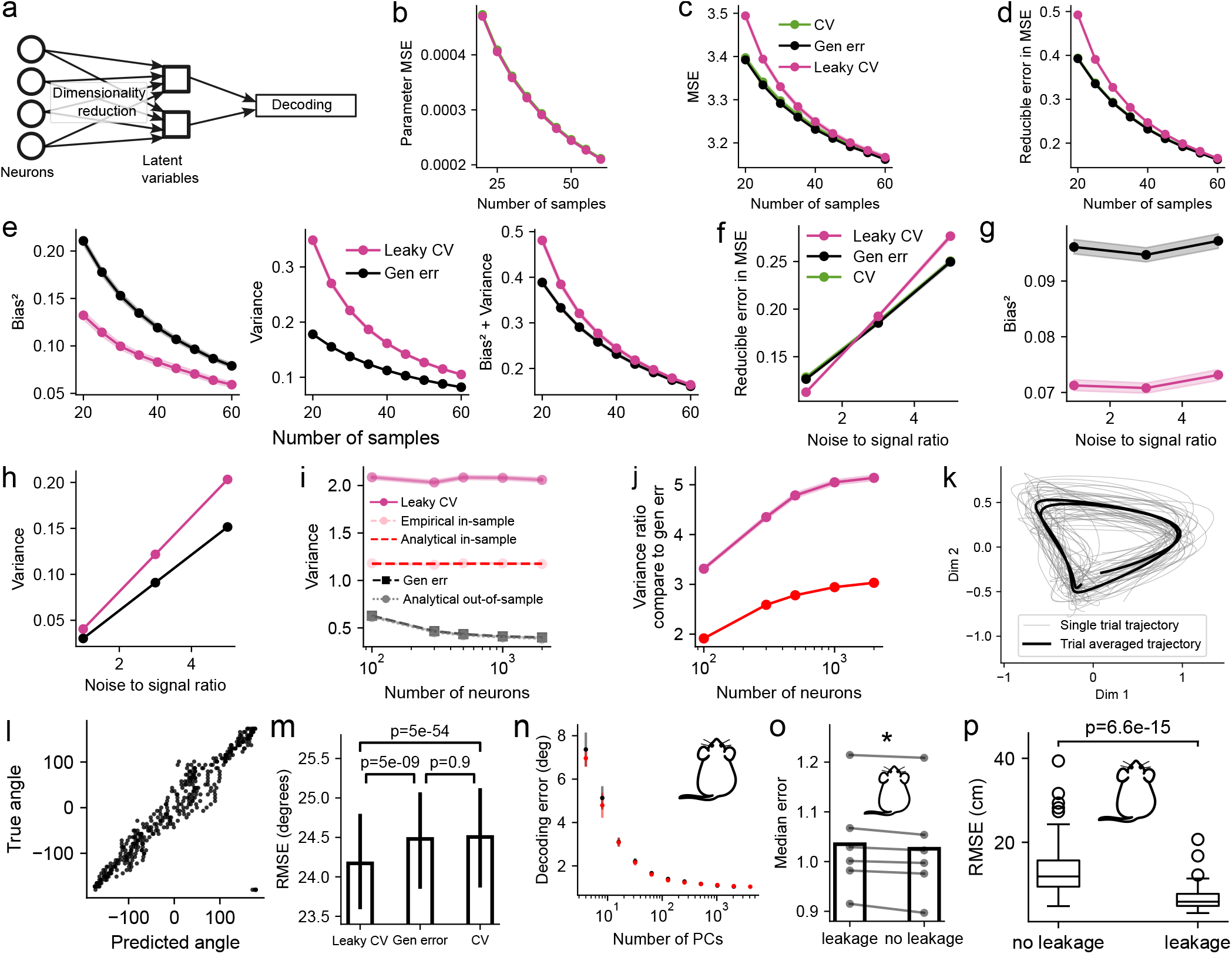
Data leakage through unsupervised dimensionality reduction. **(a)** Schematic of a common analysis pipeline where high-dimensional neural activity is first reduced to a low-dimensional latent space before being used for decoding. **(b-i)** With simulated data and ground truth parameters, PCR was used for unsupervised dimensionality reduction. **(b)** The MSE between the estimated parameters when compared to the ground truth parameters between the CV (non-leaky) and leaky CV case were similar. **(c)** The sample MSE showed a large difference. CV: non-leaky pipeline evaluated on the held-out point ***x***_0_. Leaky CV: leaky pipeline evaluated on the leaked point ***x***_0_. Gen err: leaky pipeline evaluated on many new points ***x***_new_. **(d)** Same as (c) but with the irreducible error removed in the MSE. Reducible error in MSE = Bias^2^ + Variance. **(e)** Bias-variance decomposition of a PCR decoder’s error as a function of sample size. The estimated bias term is consistently underestimated (optimistic), while the variance term is overestimated (pessimistic). This demonstrates that the total error from leakage when compared to true generalization error can be biased in either direction, depending on the noise level which scales the variance term. Here the eigenspectrum of the signal was assumed to be power-law. **(f)** Varying the ratio between signal and noise strength, the MSE in the leaky pipeline can become higher or lower than its generalization error or non-leaky CV pipeline. **(g)** Bias^2^ term in (f). **(h)** Variance term in (f). **(i)** Focusing on the variance term and varying the number of neurons in the simulation, the analytical out-of-sample term closely approximates the generalization error. Empirically, the leaky CV estimate can be bounded from below by the in-sample error (both empirical and analytical). **(j)** The variance term can be increasingly overestimated as the number of neurons (features) increases. (continued) **(k**,**l)** Simulation using GPFA for dimensionality reduction prior to decoding. **(k)** A 2D latent variable representing ground truth stimulus angle. Single-trial (gray) and trial-averaged (black) latent trajectories recovered from simulated neural activity using GPFA. **(l)** A k-nearest neighbors decoder accurately predicts the true angle from the recovered latent variables on held-out test data. Each dot represents one held-out time-bin sample at a different angle. **(m)** Comparison of decoding errors from three pipelines. The cross-validated error from a leaky pipeline (*Leaky CV*) is significantly different from both a correctly implemented, *non-leaky CV* and the true generalization error of the leaky model (*Gen error*). Wilcoxon signed-rank test, n=1000 runs. **(n**,**o)** Application to real neural data from Stringer et al. (2021). **(n)** The decoding error from a model with leaky preprocessing (red) shows a slight optimistic bias compared to the non-leaky model that uses within-fold preprocessing (black). Error bars denote SEM across animals. The rodent icons indicate that panels {n,o,p} use data from rodent experiments (mouse: panels {n,o}; rat: panel p). **(o)** Correcting data leakage in just the z-scoring step of a decoding pipeline consistently reduces the median decoding error. Each connected pair of dots represents one animal. Wilcoxon signed-rank test. **(p)** Unsupervised CEBRA (Schneider et al., 2023) was used on data same as Fig. 3f. Decoding accuracy is inflated in a pipeline with data leakage compared to a pipeline without leakage. Wilcoxon signed-rank test.

Given that the variance term drives the pessimistic behavior of PCR under leakage (see Sections S4.4 to S4.8 for detailed derivations and comparisons of the variance term in different cases), we next examined its scaling with dimensionality (see Section S4.9 for the simulation setup across different dimensions). We observed that the leaky pipeline’s variance term was bounded below by the in-sample variance (derivations in Section S4.8), which remained constant across dimensions (Fig. 5i). The variance entering the true generalization error was closely approximated by an out-of-sample estimate (Fig. 5i). Both the leaky and in-sample variance terms could be multiple-fold larger than the generalization variance term (Fig. 5j). Random matrix theory results show that out-of-sample variance can exceed or fall below in-sample variance (Green & Romanov, 2025), providing theoretical support for overestimation of the variance term in the leaky case. Taken together, PCR is reliable only when PCA is refit within each fold; globally fitting PCA miscalibrates CV errors and should be avoided.

We then used simulations to illustrate how unsupervised dimensionality reduction (here Gaussian Process Factor Analysis (GPFA)) as a preprocessing step can alter decoding accuracy when the test trials are included during dimensionality reduction. Neural spikes were simulated from a two-dimensional latent angular variable, and GPFA recovered 2D latent trajectories (Fig. 5k) from the binned firing rate across many simulated trials. Such trajectories were then used with a k-nearest-neighbor regressor (k=5) to decode angle (Fig. 5l). Fitting GPFA globally before CV leaked test information, producing over-optimistic decoding error relative to an approximate generalization error (Fig. 5m). In contrast, fold-wise refitting removed leakage and yielded CV errors that matched generalization error (Fig. 5m).

To demonstrate these effects on real neural data, we re-analyzed two population recordings from visual cortex of awake mice (Stringer et al., 2021). In the first dataset, the original pipeline z-scored and applied PCA on the full dataset. Refitting both steps within each CV fold produced a small but consistent increase in decoding error, revealing that a leaky pipeline where z-scoring and PCA were applied globally optimistically underestimated error (Fig. 5n), with sign consistent with our PCR bias–variance analysis in the high SNR regime (Fig. 5e,f). In the second dataset, for which the original pipeline (Fig. 2H of Stringer et al. (2021)) used z-scoring as the only preprocessing step, correcting this single source of leakage consistently reduced median decoding error across animals (Fig. 5o). Because the original recordings contained thousands of stimuli, these corrections changed the error estimates only modestly and did not alter the qualitative conclusions of the original study. However, when we downsampled the same recordings to tens or hundreds of stimuli, a regime closer to many other neural decoding experiments, the difference between leaky and fold-wise preprocessing became substantially larger (Fig. S7). Similar to previous analysis on decoding the animal location (Fig. 3f), we then used unsupervised CEBRA (Schneider et al., 2023) for dimensionality reduction, and again we observed that the leaky pipeline optimistically underestimated error (Fig. 5p). This confirmed that data leakage via unsupervised preprocessing can unpredictably skew model evaluation in real-world neuroscience applications, reinforcing the need for correctly implemented, within-fold preprocessing to ensure valid and reproducible results.

### 2.4 Data leakage from temporal correlations in neural time-series

Beyond preprocessing, data leakage can also arise from how samples are split for CV. CV assumes that held-out samples are independent of the samples used for training, however neural time series often violate this condition. Autocorrelation can arise, for instance, from the slow biophysical kinetics of calcium indicators, population-level synchronous dynamics reflected in LFP and EEG signals, refractory periods, or slowly varying internal states (Harris & Thiele, 2011; Cowley et al., 2020). Such temporal structure can affect decoding analyses in two distinct ways. First, shared slow temporal structure between task variables and neural activity can produce apparent decoding in the absence of genuine task encoding. The pseudosession framework of Harris (2021) addresses this problem by constructing empirical null distributions that test whether neural activity contains task information not explained by design-related temporal structure. Second, when neural activity carries task information, randomized splitting can underestimate decoding error by placing temporally adjacent samples in both training and test sets.

To address this second failure mode, we simulated calcium-imaging data with within-trial temporal correlations but no across-trial correlations. Spiking activity was generated from neuronal tuning curves encoding an angular variable, then convolved with a calcium-indicator kernel, producing autocorrelated fluorescence traces (Fig. 6a). Randomized k-fold CV (randomized CV) split individual time points across training and test sets, reflecting the statistical coupling that arises whenever random splitting places temporally dependent samples from the same trial into different folds (Fig. 6b). By contrast, leave-one-group-out CV (LOGO CV) held out entire independent trials, ensuring that temporally adjacent samples from a single trial were not split across folds (Fig. 6c). In both schemes, preprocessing and hyperparameter selection were restricted to the training data. Therefore, any mismatch between cross-validated and generalization error reflected temporal dependence induced by the split rather than data leakage from model selection or preprocessing. LOGO CV estimates more closely matched generalization error than randomized CV, indicating that randomized CV produced optimistic error estimates (Fig. 6d).

**Figure 6:**
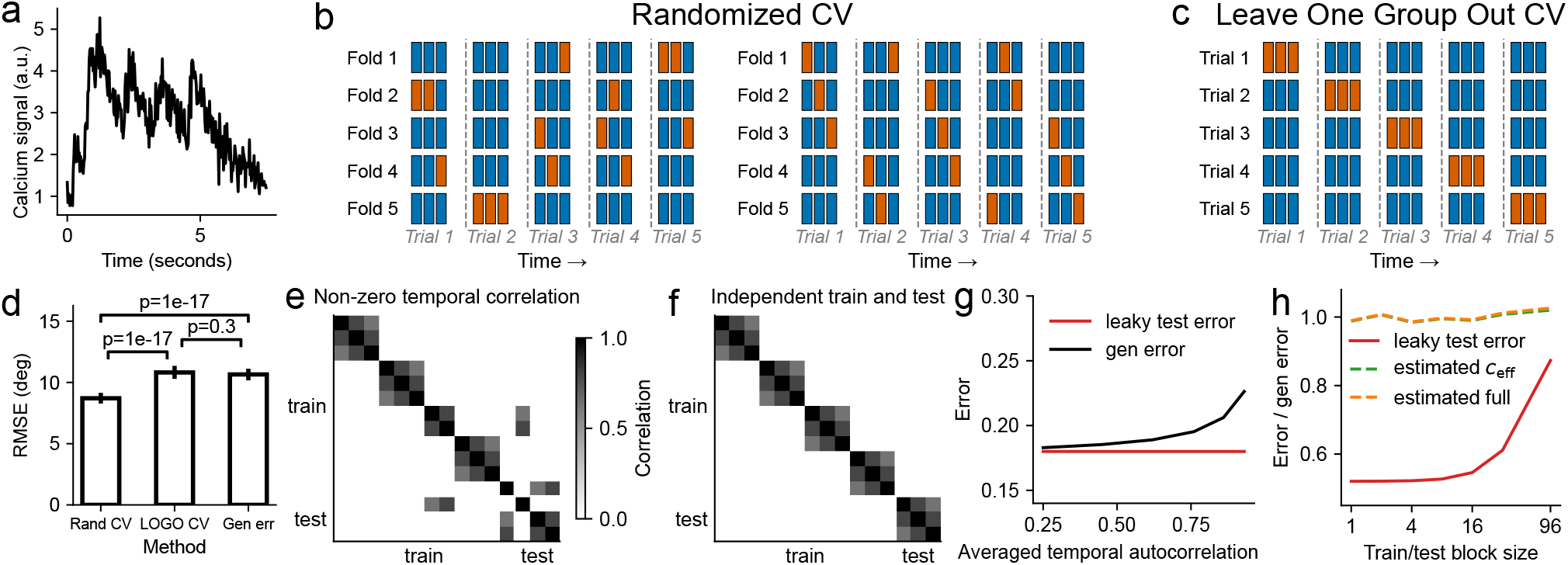
Data leakage from inappropriate CV of autocorrelated neural time series. **(a)**: Example of a continuous, autocorrelated neural signal, such as that found in EEG, LFP, or calcium imaging data (simulated calcium trace shown here). The signal’s value at any given time point is highly correlated with its value at adjacent time points. **(b)**: Two flawed validation schemes using standard randomized k-fold CV (randomized CV). Data points were assigned to training (blue) and test (orange) sets randomly, which violated their temporal structure and trial-based independence. **(c)**: Given the trial-based independence in the simulation, a more methodologically sound approach using leave-one-group(trial)-out CV (LOGO CV). Entire independent trials were held out as the test set. This correctly preserved the temporal structure within each trial while ensuring no temporal dependence existed between the training and test data. For both methods, hyperparameter optimization was performed using a nested CV loop (omitted from illustration for clarity). **(d)**: Randomized CV led to artificially inflated performance metrics. The decoder evaluated with the randomized CV (“leaky”) method showed a significantly lower RMSE than one evaluated with LOGO CV (“non-leaky”) However, the true generalization error, assessed on a separate, held-out test set, revealed that the leaky method’s superior performance was an artifact. LOGO CV, in contrast, provided an accurate estimate of how the model would perform on new, unseen data. **(e)**: Covariance matrix of the simulated signal from Fold 1 under randomized CV setting in (b), reordered so that training samples precede test samples on both axes. Within-trial autocorrelation produced non-zero entries in the off-diagonal train × test block, reflecting statistical dependence between the training and test sets that inflated the apparent decoding performance. The second scheme in (b) has higher averaged temporal correlation than the first scheme. **(f)**: The same covariance reordered from any single fold under LOGO CV in (c). Because the held-out trial shared no samples with any training trial and cross-trial correlations were zero, the train × test block vanished, indicating statistically independent training and test sets. **(g)**: In a simplified binary-decoding setting, the same apparent leaky test error corresponded to different true generalization errors as the train–test temporal correlation varied. Lower values along the horizontal axis indicate weaker train–test correlation, whereas higher values indicate stronger correlation. **(h)**: Covariance-based corrections brought the leaky estimate closer to performance on independent data across train/test block sizes; the estimated corrections were closer to the true generalization error than the naive leaky test estimate. The scalar correction *c*_eff_ collapsed the per-sample train–test covariance reductions to their class-averaged mean and applied a closed-form variance adjustment; the full correction instead retained the per-sample leakage vector and numerically inverted the test-error equation to recover the generalization variance (see Methods and Section S5). The block size determined the length of contiguous segments alternately assigned to the training and test sets: smaller blocks placed more training and test samples adjacent in time, producing stronger train–test correlation, whereas larger blocks separated them further and reduced this correlation.

The covariance structure makes the source of the bias explicit. Under randomized CV, within-trial autocorrelation produced non-zero train *×* test entries in the covariance matrix, showing that held-out samples were statistically coupled to training samples (Fig. 6e). Under LOGO CV, held-out samples came from an independent trial. Because cross-trial correlations were zero in the simulation, the train *×* test block vanished (Fig. 6f). Thus, leakage through temporal correlation can persist even when all fitting operations are confined to training data, because the split itself determines whether the test set is independent.

In a simplified binary-discrimination task, we then tested whether train–test covariance was sufficient to explain the error-estimation difference. The class labels of the stimuli contained no slow structure and were independent of time, so the difference only arose because autocorrelated, task-irrelevant noise made held-out samples statistically predictable from nearby training samples. An analytical calculation for a linear decoder showed that the relevant quantity is the average covariance between each held-out sample and the training samples used to estimate the decision boundary (see Methods). Specifically, while the generalization error remained determined by the true class separation, strong train–test covariance could make apparent leaky test error lower than the true generalization error. Consistent with this, when class separation was adjusted to hold the apparent leaky test error fixed, increasing train–test temporal correlation produced different true generalization errors (Fig. 6g).

We then asked whether this covariance structure could be used as a practical diagnostic. We varied the size of contiguous train/test blocks, which changes the train–test covariance while leaving the marginal noise model and decoder fixed. Smaller blocks placed more training and test samples close together in time, whereas larger blocks separated them more strongly. For each split, we estimated temporal covariance from residual activity and used the train–test covariance block to correct the leaky test error toward true generalization error. Across block sizes, the corrected estimates were closer to true generalization error than the uncorrected leaky estimate (Fig. 6h). Together, these analyses identify train–test temporal covariance as a measurable and correctable source of optimistic error estimation in autocorrelated neural time series.

## 3 Discussion

Here we systematically examined how data leakage can arise in common neural decoding pipelines and how it biases empirical performance estimates. The empirical analyses are complemented by derivations that explain why these biases arise. Using a range of supervised and unsupervised preprocessing methods, we showed that fitting preprocessing steps on the entire dataset before CV can lead to optimistic or pessimistic errors that do not reflect true generalization performance. Across simulations and real datasets, we demonstrated that properly implemented train-test splits, in which all parameters of the preprocessing and decoding pipeline are learned only on the training data within each fold, restore calibration between cross-validated error and generalization error. In the Recommendations section (Section S1) we outline simple templates for leak-free neural decoding pipelines, and we propose a practical diagnostic to help detect potential leakage in existing analyses.

For temporally correlated recordings, preventing leakage also requires attention to the unit over which independence is approximated. Preprocessing only within each fold is necessary, but it does not ensure that held-out samples are independent if nearby time points are split across training and test sets. Whenever possible, validation should hold out experimental units more likely to be independent, such as complete trials or sufficiently separated blocks. These units should be chosen to match the scientific quantity to be estimated. For within-experiment decoding, the goal is usually to evaluate held-out samples from the same experimental context while minimizing train-test dependence. When no natural independent unit is available, block-size sensitivity analyses and covariance-based diagnostics can help assess whether the reported decoding error depends on residual train-test covariance. However, these analyses should be treated as diagnostics rather than substitutes for a well-designed validation split.

Our focus has been on preventing data leakage when training and evaluation data are intended to represent the same target distribution. This is distinct from out-of-distribution generalization, which asks whether a decoder transfers across recording days, subjects, tasks, or other experimental contexts. A decoder may be well calibrated for appropriately held-out trials within one experiment yet fail in a new context, and such failure reflects limited generalization rather than data leakage. Researchers should be explicit about whether their conclusions concern information content within a fixed experimental context, or claims about robustness across contexts.

Current intracortical BCI research actively targets this generalization challenge. To maintain a fixed decoder despite recording instabilities, methods align latent dynamics or full-dimensional neural distributions across recording days (Degenhart et al., 2020; Karpowicz et al., 2025; Ma et al., 2023). However these alignment techniques are themselves data-dependent, even when unsupervised. Thus hyperparameter selection, alignment method choice, and any representation update must be validated without access to the held-out evaluation day.

The emergence of large-scale neural foundation models trained on extensive datasets from multiple subjects and experimental conditions (e.g. Ye et al. (2023); Azabou et al. (2023, 2025); Wang et al. (2025)) presents both opportunities and challenges for preventing data leakage. These models provide pre-trained representations that capture generalizable neural patterns across diverse experimental contexts and subjects. To apply these models to new data, researchers typically need to adapt the pre-trained representations through self-supervised (or supervised) learning. These adaptation steps can include hyperparameter selection, normalization, alignment, and test-time adaptation. Each is effectively another form of preprocessing, where the model learns to map new neural data into the pre-trained representation space. Pretraining on external data does not itself introduce leakage, provided the held-out test set is excluded from the representation update. However, if the foundation model is adapted using the entire dataset before splitting into training and test sets, information from the test set will contaminate the representation learning process. For claims about held-out samples within one recording, adaptation steps must be fit without those held-out samples. For transfer claims across recording days, animals, subjects, or tasks, the evaluation unit being tested should remain outside the adaptation process. The only exception is when test-time adaptation is itself the deployment setting; in that case, the adaptation rule should be specified in advance and assessed on a further, completely unseen dataset.

The same principles extend beyond neural decoding. Recent work on alignment between large language models and the brain showed that shuffled train-test splits and overlooked confounds can create apparent brain-model correspondence that is not robust under stricter evaluation (Hadidi et al., 2026). As these pipelines grow in complexity, so do the opportunities for data leakage, making it essential to verify that every data-dependent step is isolated from the held-out data for evaluation.

## 4 Methods

### 4.1 Zebrafish

All procedures were performed with approval by the Animal Studies Committee at Washington University and adhere to NIH guidelines. Nacre zebrafish (*Danio rerio*) embryos expressing elavl3:H2B-GCaMP6s were raised from birth to 8/9 days post-fertilization under 14/10 light/dark cycles at 28.5°C. 3 h prior to imaging, zebrafish larvae were embedded in 2.5% low-melting point agarose in the center of 35 mm diameter Petri dishes immersed in E3 embryo medium. 2-photon images were acquired using a Bruker Ultima 2pPlus microscope with a 16X/0.8NA water-dipping lens (Nikon). Samples were excited via a Coherent Discovery TPC-one tunable laser at 940nm. Volumetric images consisting of 10 planes were taken starting from 30 µm to 120 µm from the dorsal surface in 10 µm steps at a rate of 2.1Hz/volume. Visual stimuli were projected onto white diffusion paper placed in front of the chamber using a pico-projector (PK320 Optoma, USA). Larvae were placed in the center of the chamber with one eye facing the projected side of the dish with body axis at 90°to the projector direction. Visual stimuli were generated using custom software based on MATLAB (MathWorks). Dark spots of 6°diameter at 11 different azimuth locations from 15°to 165°in 15°intervals were presented to the right visual field, with 0° being the body axis of the fish. Stimuli were presented in 10 blocks of the 11 stimuli, where stimuli within each block were of different order. Each stimulus lasted 1 second and a stimulus was presented every 10 seconds. Neural recordings were motion-corrected and converted to Δ*F/F*_0_ traces using CaImAn (Giovannucci et al., 2019). The activity evoked by a single stimulus was defined as the averaged calcium activity of the third and fourth frames after stimulus onset.

### 4.2 Gaussian activity (Fig. 2)

We simulated two multivariate Gaussian distributions with distinct means and identical covariance matrices: *N* (***µ***_1_, ***I***_*p×p*_) and *N* (***µ***_2_, ***I***_*p×p*_). Here, ***µ***_1_ ∈ ℝ^*p×*1^, ***µ***_2_ ∈ ℝ^*p×*1^, where *p* represented the number of neurons in the simulation. ***µ***_1_ and ***µ***_2_ were randomly generated according to the tuning profile of the neurons, while keeping ∥***µ***_1_ − ***µ***_2_∥_2_ the same.

The optimal discriminant function had an overall error rate equal to 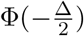 where Φ(·) denotes the cumulative probability distribution for the standard Gaussian, and Δ is the Mahalanobis distance 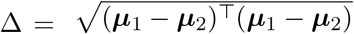 (Mahalanobis, 1936; McLachlan, 1999). In our simulations, Δ was assigned to unity, resulting in an error rate of approximately 0.31 and an optimal accuracy of 0.69.

We drew 300 random samples from each distribution to represent neural activity in response to stimuli, simulating the number of trials in a typical neuroscience experiment.

### 4.3 Projecting neural activity to PC space defined by PCA on trial-averaged data (Fig. 3)

We generated random neural activity where each neuron had a random response simulated independently to stimuli. Random labels for each population vector were also generated from the 11 stimulus locations (15°to 165°in 15°steps), each with 20 repetitions. 2000 neurons were simulated, resulting in a 20×11×2000 matrix. PCA was then performed on the trial-averaged matrix, and the singular vectors corresponding to the k largest singular values were used for projecting the high-dimensional neural activity to a low-dimensional subspace. We used k=2 for visualization and k=3 in the relevant decoding pipeline. Different data (all trials or trial-averaged) were projected into this space (Fig. 3a,d).

### 4.4 Simulation of Neural Activity (Fig. 4)

We simulated neural population responses to a stimulus variable, *θ*, representing an angle from 0 to *π*. On each trial *t*, the stimulus was drawn from a uniform distribution, *θ*_*t*_ ~ *U*(0, *π*), and represented as a 2D vector ***x***_*t*_ = [cos(*θ*_*t*_), sin(*θ*_*t*_)]^⊤^.

The activity of a population of *N*_neurons_ was simulated using two variations of a Gamma-Poisson generative model, which captured the Poisson-like statistics and overdispersion commonly observed in neural data.

#### 1. Parametric Gamma-Poisson (Gaussian Tuning)

Each neuron *i* was assigned a fixed preferred stimulus angle, *µ*_*i*_ ~ *U*(0, *π*). A deterministic mean firing rate, *f*_*i*_(*θ*_*t*_), was defined as

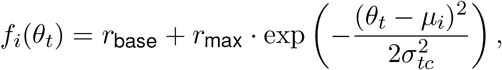

where *r*_base_ is a baseline firing rate, *r*_max_ is the maximum rate above baseline, and *σ*_*tc*_ is the tuning width. *λ*_*i,t*_ was then drawn from a Gamma distribution,

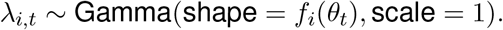

#### 2. Semi-Parametric Gamma-Poisson (Linear Projection)

To use a more flexible, non-parametric tuning assumption, a linear projection 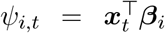 was constructed, where ***β***_*i*_ was drawn from a standard normal distribution. The corresponding rate parameter was defined as 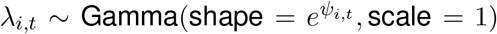. A subset of neurons was designated as “high-rate” by multiplying their generated *λ*_*i,t*_ values by a constant factor.

For both models, the final spike count was drawn from a Poisson distribution, *y*_*i,t*_ ~ Poisson(*λ*_*i,t*_).

For each simulation run, a single “master” dataset of neural responses was generated for the maximum number of samples, along with a large, independent held-out test set (10,000 samples) for estimating generalization error. We then iterated through a range of smaller sample sizes, *n*, by taking the first *n* trials from the master dataset. Unsupervised feature selection was performed by selecting the top *k* neurons with the highest variance. The exact model from the leaky procedure was evaluated on the 10,000-sample held-out test set to compare the estimated decoding error from CV and the true generalization error.

This entire process was repeated for 10,000 independent simulation runs to generate stable estimates and error bars for each metric as a function of the number of samples. For the neuron-count sweep used to contextualize this effect, we used the parametric Gamma-Poisson model with Gaussian tuning curves, fixed the total number of samples to 100, used 5-fold cross-validation, selected the top 10% of neurons by evoked spike-count variance, and varied the total number of recorded neurons up to 750 so that the selected feature count remained below the number of training samples per fold.

### 4.5 PCR Simulation (Fig. 5)

We considered a standard linear model where the response *y* was generated from *d*-dimensional features ***x*** with a ground truth coefficient vector ***β*** and additive noise 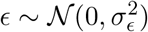:

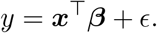

The features ***x*** were drawn from a multivariate normal distribution *N* (**0, Σ**). In our simulations, the signal vector ***β*** was constructed to lie within the subspace spanned by the top *k* eigenvectors of the true population covariance **Σ**. Data generated from this model were also used in PLS regression in Fig. 3.

#### 4.5.1 The Leaky Procedure and Error Metrics

Our simulation framework evaluated the leaky PCR procedure in which held-out data contributed to preprocessing (PCA) before regression, and compared its apparent test error with its true generalization error. The notation below uses a representative held-out point ***x***_0_, matching the implementation in Fig. 5. Detailed derivations for the pointwise formulation below and for the corresponding multiple-point extension are given in Section S4.

For a fixed training set of *n* samples, (***X***_train_, ***y***_train_), and a random point ***x***_0_ drawn from the same feature distribution, a “leaky” PCA basis 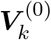 was computed from the augmented dataset that included ***x***_0_. A linear regression model was then trained using only the original training data projected onto this leaky basis. This procedure produced a different model, 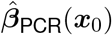, for each realization of ***x***_0_.

We then compared two error estimates for this leaky model:

1. **The Leaky Estimate:** the error when the model was evaluated on the specific point ***x***_0_ that was used to create the leaky basis. This mimicked the biased estimate one might obtain when test points influenced the preprocessing step.
2. **The True Generalization Error:** the error of the exact same model when evaluated on a new, unseen data point ***x***_new_, drawn from the true underlying distribution. This corresponded to the ideal generalization performance of the leaky model.

#### 4.5.2 Bias-Variance Decomposition and Comparison

To understand how leakage affects performance, we decomposed the Mean Squared Error (MSE) for both estimates into bias and variance components. The expected prediction error for a model predicting on a point ***x*** is:

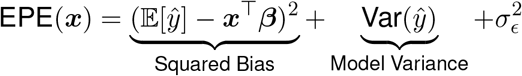

The expectations for our two error estimates are taken over the distribution of ***x***_0_, as it defines the model. For the *Leaky Estimate*, evaluated at ***x***_0_:

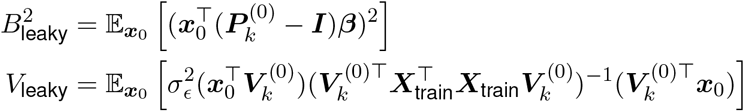

For the *True Generalization Error* of the same leaky model, evaluated at a new point ***x***_new_:

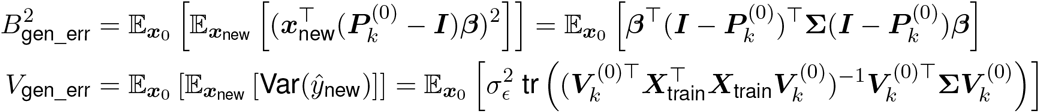

where 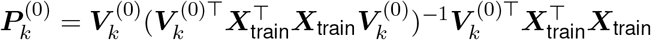.

#### 4.5.3 Monte Carlo Estimation

A direct analytical comparison of the leaky terms versus their true generalization error counterparts was intractable, as the expectations over ***x***_0_ involved complex dependencies on the eigenvectors of a randomly perturbed matrix. Therefore, we estimated these expected values using Monte Carlo simulations. For each independent simulation run, we generated a fixed training set ***X***_train_ where each ***x***_*i*_ ~ *N* (**0, Σ**) and then, for that fixed set, averaged the bias and variance terms over many draws of ***x***_0_. This provided a stable and accurate numerical estimate of the expected terms for comparison.

We wrote **Σ** = ***Q*Λ*Q***^⊤^ with two different forms of eigenspectrum (diagonal elements of **Λ**) used in Fig. 5: (i) a spiked covariance matrix (Fig. S5) and (ii) a smoothly decaying spectrum (Fig. 5). The eigenspectra were normalized to *d* such that tr(**Λ**) = *d*. Orthogonal matrices ***Q*** were drawn randomly from the Haar distribution.

### 4.6 Neural activity encoding heading angle (Fig. 5)

Following Denker et al. (2018), we simulated an animal continuously rotating its heading angle *θ*(*t*) at a constant angular speed, with each full rotation (trial) lasting 2 seconds. The two-dimensional activity representing the heading angle was simulated as [sin (*θ* (*t*)), cos (*θ* (*t*))] ^⊤^, which we considered the ground truth latent activity. The 2D latent activity was projected into a 50-dimensional space using a random matrix ***P*** ∈ ℝ^50*×*2^. This activity was normalized and exponentially transformed to ensure non-negativity of the firing rates, with baseline firing rates added as noise. Spikes were then generated using an inhomogeneous Poisson process. We simulated 20 trials in total. GPFA (Yu et al., 2009) was then used to solve the inverse problem of recovering the matrix ***P*** and the 2D latent activity from the observed spikes.

### 4.7 Calcium activity simulation (Fig. 6)

To study the effect of autocorrelation on CV, we generated synthetic calcium fluorescence data following the commonly-used generative model of an autoregressive process (Vogelstein et al., 2010; Pnevmatikakis et al., 2016; Friedrich et al., 2017). For each neuron, spikes *s*_*t*_ were drawn from a Poisson process, and calcium evolved as *c*_*t*_ = *γc*_*t*−1_ + *s*_*t*_, with *γ* = exp(− Δ*t/τ*). The observed fluorescence was *y*_*t*_ = *c*_*t*_ + *b* + *ϵ*_*t*_, with non-negative baseline *b* and measurement noise *ϵ*_*t*_ ~ *N*(0, *σ*^2^). We used an AR(1) model at 40 Hz and *τ*=1.24 s. We simulated N=50 neurons, organized into trials by resetting *c*_0_ at the start of each trial. Each trial lasted 300 frames. 20 trials were simulated and used for CV, and an additional 10 trials were simulated for evaluating generalization error. We compared two CV schemes: (i) randomized *k*-fold CV that split individual time points at random; (ii) leave-one-trial-out CV, holding out entire trials. For both schemes, any preprocessing was fit on the training data only and then applied unchanged to the held-out data. Neural activity and stimulus labels were simulated similar to Section 4.6. Nested CV was used for PCR to select the number of PCs, and the selected model was evaluated on the held-out test set.

### 4.8 Temporal-covariance leakage simulations and correction (Fig. 6)

For the simplified binary-discrimination simulations in Fig. 6g,h, we considered balanced positive and negative classes with scalar responses drawn from a multivariate Gaussian whose temporal covariance followed an exponential kernel, Σ_*ij*_ = *ρ*^*λ*|*i*−*j*|^ (equivalently Σ_*ij*_ = *α*^|*i*−*j*|^ with *α* = *ρ*^*λ*^ and unit marginal variance; Section S5 writes the same kernel as *σ*^2^*α*^|*i*−*j*|^). A one-dimensional LDA classifier was trained on the training partition, and its test error was compared to the true generalization error on independent data. We analytically derived the relationship between the train–test covariance block and the resulting bias in the test error estimate (full derivations in Section S5).

For Fig. 6g, we used 96 time points with an alternating one-sample train/test split, fixed *λ* = 0.55, and varied *ρ* ∈ {0.25, 0.45, 0.62, 0.76, 0.86, 0.93} while tuning the signal mean *µ* so that the analytical leaky test error remained fixed at 0.18. For Fig. 6h, we fixed *ρ* = 0.94 and *λ* = 0.5, varied alternating contiguous block sizes {1, 2, 4, 8, 16, 48}, and compared the uncorrected leaky error with two covariance-corrected estimates. The scalar correction (“estimated *c*_eff_”) collapsed the per-sample train–test covariance reductions to their class-averaged mean and applied a closed-form variance adjustment. The full correction (“estimated full”) retained the per-sample leakage vector and numerically inverted the test-error equation to recover the generalization variance, accounting for heterogeneous leakage across held-out samples. Both corrections used a temporal covariance estimated from residual activity via a semivariogram-based fit rather than the ground-truth covariance, and were averaged over 600 covariance-estimation replicates, each using 16 calibration residual traces. Full mathematical details of both correction procedures and the covariance estimation are given in Section S5.

## 5 Data & code availability

The source data and code will be made publicly available upon publication.

## Supplementary Information

### S1 Recommendations

#### S1.1 General pipeline guidelines

A recommended pipeline with cross-validation to assess decoding accuracy avoiding data leakage is described below. A Python code template for this pipeline will be made publicly available upon publication.

Step 1. Record neural activity and stimulus/behavior labels.

Step 2. Split to training and test sets.

Step 3. Apply preprocessing, feature selection and/or dimensionality reduction to the training set (optionally with labels) as necessary.

Step 4. Train the decoder on the processed training set.

Step 5. Apply the same preprocessing steps and transformations as learned in Step 3 to the test set.

Step 6. Evaluate the decoder and report performance on the processed test set.

Step 7. If cross-validation is used, repeat Step 3–6 with the new split.

The pseudocode provided in Algorithm 1 illustrates this pipeline in more detail. Additionally, Figure 1f,g offer a visual comparison of both pipelines, highlighting the critical steps where the recommended approach diverges to maintain the integrity of the validation process.

##### Algorithm 1

A recommended pipeline with cross-validation to assess decoding accuracy avoiding data leakage

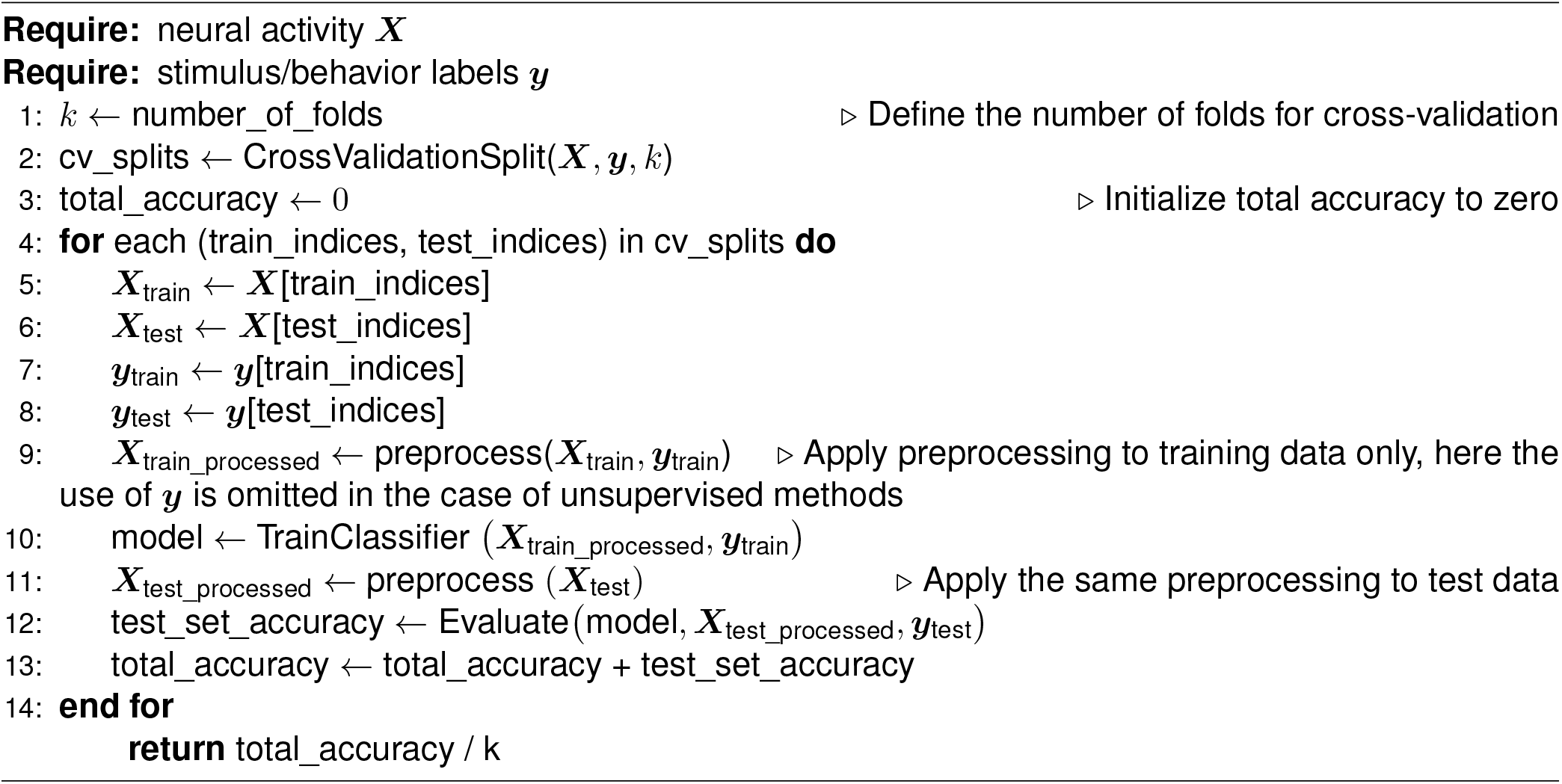

To assess decoding accuracy for temporally correlated data, see Section 2.4 and the Discussion section.

#### S1.2 Detection methods

We suggest a simple practical check to test for data leakage. An effective approach for determining the chance level in neural decoding analysis is through permutation tests, also known as shuffle controls (Ojala & Garriga, 2010; Combrisson & Jerbi, 2015). This method involves permuting stimulus identities or neural activity across classes to establish an empirical null distribution of decoding accuracy, providing a more accurate estimate than the naive 1/(number of unique stimuli) calculation. For a balanced classification problem, if a shuffle control is performed on an entire decoding pipeline which involves data leakage, the observed null distribution will be different than the 1/(number of unique stimuli) estimate.

However in typical neuroscience analyses, a shuffle control is performed just before decoding. The null distribution constructed this way does not incorporate preprocessing steps, making it less informative than a null distribution constructed by performing the shuffle control at the start of the pipeline. We therefore recommend conducting permutation tests prior to data preprocessing rather than afterwards. Since preprocessing parameters are derived from the data itself, this step should be thought of as part of the machine learning classifier. This approach is analogous to introducing randomness at the start of simulations as a null model (for example, in network science, random graphs are constructed as a null model (Barabási & Albert, 1999; Fornito et al., 2016; Váša & Mišić, 2022)). This practice also aligns with best practices in popular machine learning projects, as exemplified by the API design for scikit-learn (Buitinck et al., 2013). However note that while this check is very useful in practice, it should be viewed as a screening tool rather than a definitive guarantee: passing it reduces concern about leakage, but failing it is a clear warning sign that the pipeline needs to be audited.

#### S1.3 Hyperparameter tuning

When hyperparameter tuning is required, nested cross-validation provides a more robust approach than standard cross-validation (Stone, 1974; Cawley & Talbot, 2010). In nested cross-validation, an outer loop performs the train-test split for final evaluation, while an inner loop performs additional train-validation splits for hyperparameter optimization. This ensures that the hyperparameter selection process does not inadvertently use information from the test set, which could occur if hyperparameters are tuned on the entire dataset before cross-validation. The computational cost of nested cross-validation is higher, but it provides more reliable estimates of model performance, particularly when preprocessing steps involve hyperparameters (such as the number of components in PCA or regularization strength in many machine learning models).

### S2 PCA on trial-averaged data

#### Notation

| Symbol | Meaning |
| --- | --- |
| $d$ | Number of neurons |
| $C$ | Number of conditions |
| $T$ | Number of trials per condition |
| $k$ | Number of retained PCs ($k \leq C - 1$) |
| $\rho$ | Neuron-to-trial ratio $d/T$ |
| $S_B, S_W$ | Between-class and within-class scatter matrices |
| $J$ | Fisher discriminant ratio $\text{tr}(S_W^{-1} S_B)$ |

We derive an analytical prediction for the expected Fisher discriminant ratio 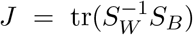 in the *k*-dimensional PC projection of a null-signal simulation, and show that *J* is governed by the ratio *d/T* of the number of neurons to the number of trials per condition.

#### S2.1 Setup

Let 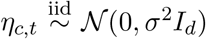 for conditions *c* = 1, …, *C* and trials *t* = 1, …, *T*. This is the null model: no true signal, random labels. Decompose each observation as

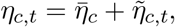

where 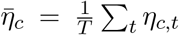 is the class mean and 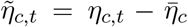 is the within-class residual. By Basu’s Theorem, 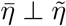, and 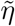 carries *C*(*T* − 1) degrees of freedom.

Trial-averaged PCA fits a truncated singular value decomposition on the *C × d* matrix 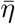 whose rows are the condition means. The top-*k* right singular vectors *W* ∈ ℝ^*d×k*^ are therefore a function of 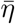 only, and are independent of 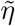.

#### S2.2 Expected Fisher ratio under the null

##### Between-class scatter

Let *H* = *I*_*C*_ − *C*^−1^**11**^⊤^ be the class-centering matrix. Under the null, the centered class-mean noise can be written as

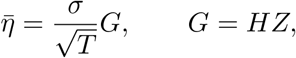

where *Z* ∈ ℝ^*C×d*^ has iid *N*(0, 1) entries. Thus *G* has rank at most *C* − 1. Let *W* contain the top-*k* right singular vectors of 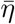, with *k* ≤ *C* − 1. Then

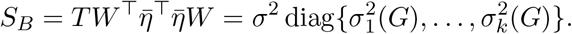

Moreover,

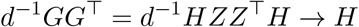

as *d* → ∞ with *C* fixed. Hence the nonzero eigenvalues of *GG*^⊤^, equivalently of *G*^⊤^*G*, satisfy

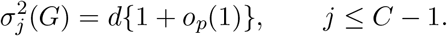

Therefore

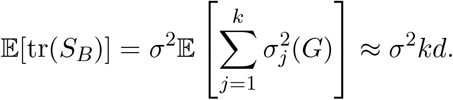

##### Within-class scatter

Since the class means and within-class residuals are independent under the Gaussian null, and since *W* is a function of the class means, conditional on *W* we have

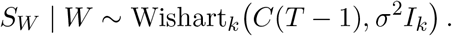

Assuming *C*(*T* − 1) *> k* + 1,

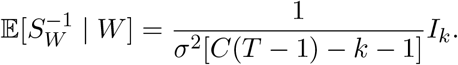

##### Combining

Using the tower rule and conditioning on 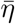, which fixes both *W* and *S*_*B*_,

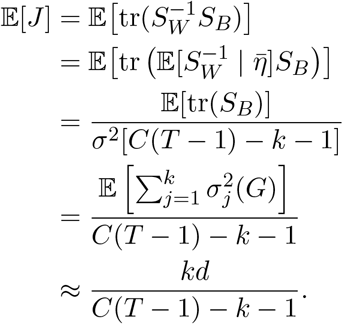

#### S2.3 *d/T* scaling

Substituting *d* = *ρT* for a fixed ratio *ρ* = *d/T*,

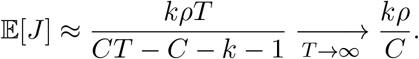

That is, *J* converges to a constant determined solely by *ρ* = *d/T, C*, and *k*. Two consequences follow:

1. At fixed *d*, increasing *T* reduces *J* approximately as *kd/*[*C*(*T* − 1) − *k* − 1] ~ *d/T*, consistent with the intuition that more repeats reduce spurious clustering (Fig. S4a).

2. At fixed *d/T, J* plateaus as *T* → ∞ (Fig. S4b). Because modern large-scale recording technology tends to scale *d* and *T* together, the pitfall does not automatically diminish with experimental scale.

##### Finite-size corrections

At small *T*, the prediction *kd/*[*C*(*T* − 1) − *k* − 1] exceeds the asymptote *kρ/C* by a factor 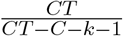, which equals ≈ 1.34 for *C* = 11, *k* = 2, and *T* = 5, and decays as *O*(1*/T*). A second finite-size correction comes from selecting the top-*k* singular directions of the centered Gaussian matrix *G*. Since *G* has rank *r* = *C* − 1, the relevant aspect ratio is 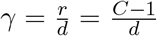. For finite *d*, the largest nonzero eigenvalues of *GG*^⊤^ exceed their limit *d*; heuristically, the upper edge is of order 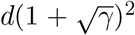.

Thus

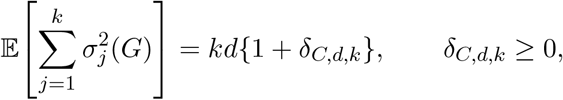

with *δ*_*C,d,k*_ → 0 as *d/*(*C*−1) → ∞. This produces an additional upward correction to the leading prediction *kd/*[*C*(*T* − 1) − *k* − 1], especially when *d* is only moderately larger than *C*.

### S3 Fixed-fraction variance selection

#### Notation

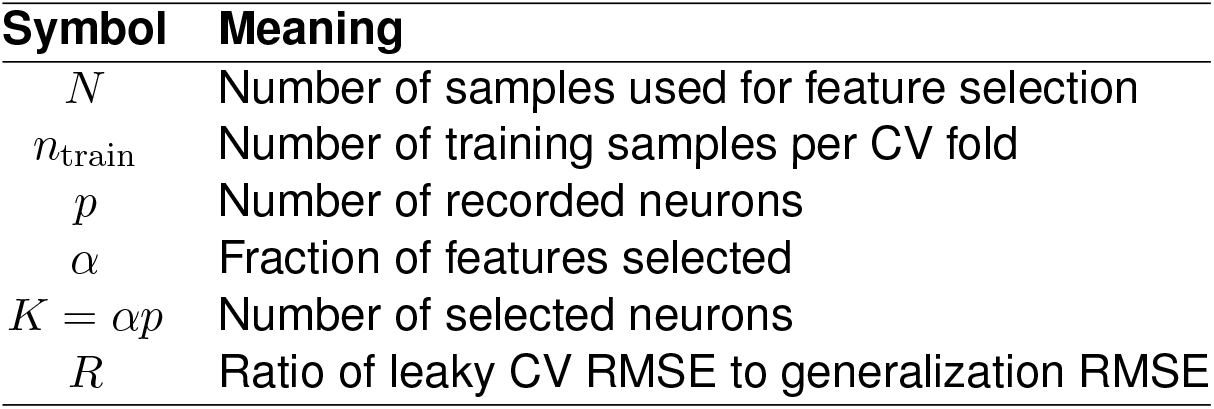

The variance-selection result in Moscovich & Rosset (2022) analyzes the extreme case in which a single feature is selected. Here we analyze the setting used in our neuron-count sweep, where a fixed fraction of features is selected and ordinary least squares remains determined. Below is not intended as an exact proof for the Gamma-Poisson spike-count simulation in the main text. Rather, it provides a benchmark on ordinary least squares (OLS) for why leakage is small in low-dimensional settings and grows as the selected feature count approaches the number of training samples.

Let *N* be the number of samples used for leaky feature selection, *n*_train_ the number of samples in each training fold, *p* the number of recorded neurons, and *K* = *αp* the number of selected neurons. Assume *K < n*_train_ − 1. For a Gaussian null feature,

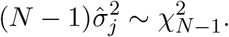

Selecting the top fixed fraction *α* does not select an increasingly extreme maximum as *p* grows. Instead, the selected features approach the upper tail of a fixed distribution. If 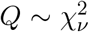 with *ν* = *N* − 1, and *q*_1−*α*_ is the (1 − *α*)-quantile of *Q*, the average variance inflation among selected features is

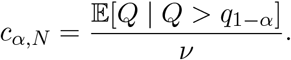

This factor is finite for fixed *α*.

For OLS with *K* selected features, the standard finite-sample variance amplification has the scale

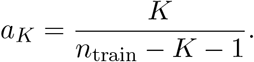

If the validation samples used in feature selection inflate the relevant selected-feature variance by *c*_*α,N*_ relative to fresh generalization samples, then a useful approximation for the ratio of leaky CV RMSE to leaky generalization RMSE is

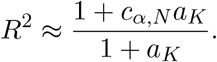

Consequently, the ratio increases with *K/*(*n*_train_ − *K* − 1) but is bounded by

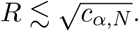

For the parameter regime used in Figure 4 *N* = 100, 5-fold CV gives *n*_train_ = 80. With *α* = 0.1 and *p* = 750, the selected feature count is *K* = 75 *< n*_train_ and

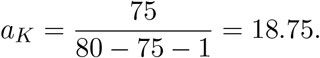

For *N* = 100 and *α* = 0.1, *c*_*α,N*_ ≈ 1.264, yielding

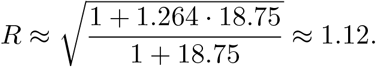

Thus, even in a stylized Gaussian setting, fixed-fraction variance selection produces little mismatch when *K* ≪ *n*_train_, but the ratio can become appreciable as *K* approaches *n*_train_ while remaining in the determined-OLS regime.

### S4 Derivations for PCR

In this section we introduce principal component regression (PCR), and derive the bias–variance decomposition of both in-sample and out-of-sample prediction risk, and we make explicit how the dependence on the PCA basis enters through an effective projection matrix *P*_*k*_(*X*_train_). The structure parallels the analysis in Green & Romanov (2025), but is adapted to our finite-sample simulation framework and to the comparison between leaky and non-leaky PCA bases.

#### Notation

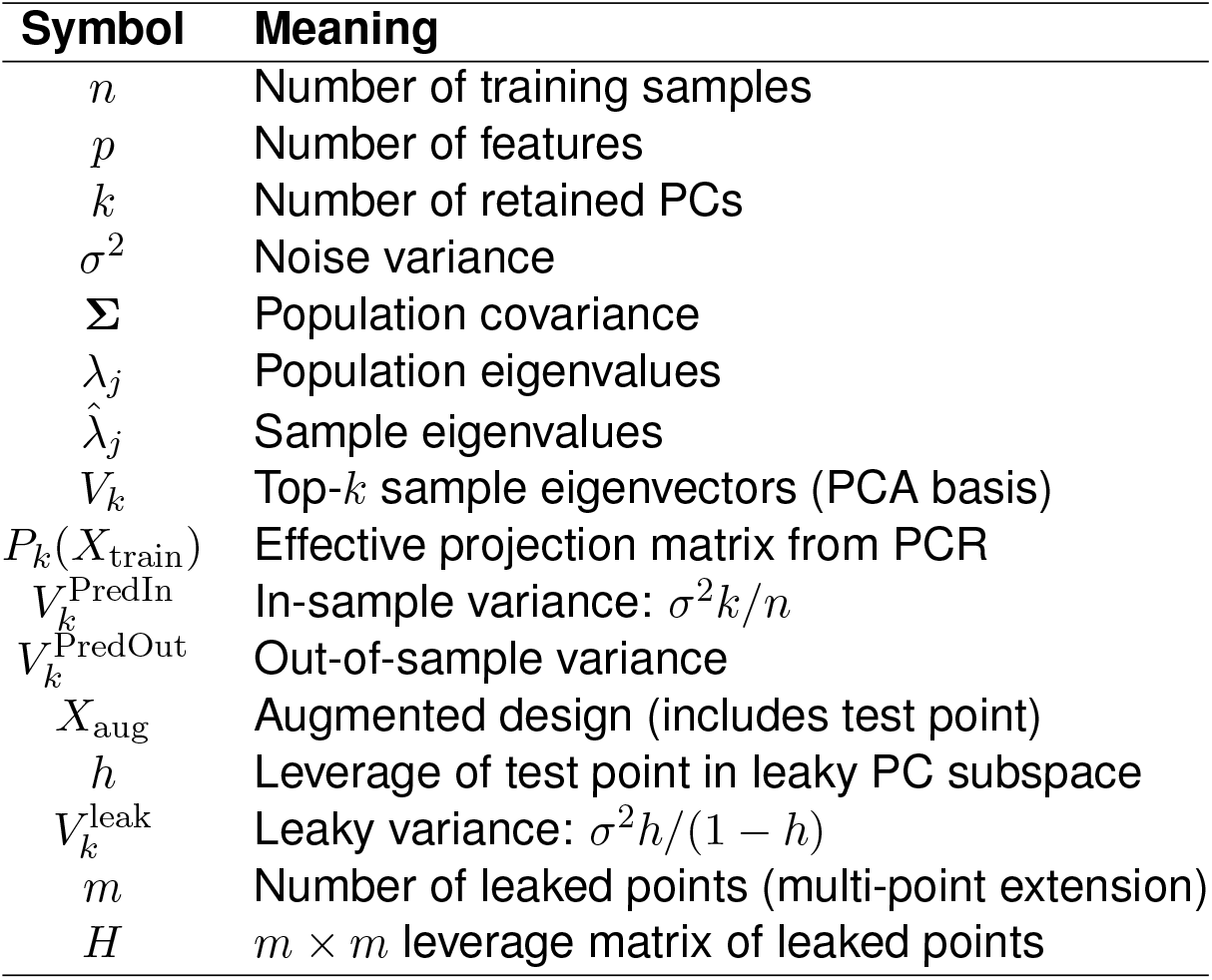

#### S4.1 Data generating process for PCR

##### Fixed number of features

Data are generated from a Gaussian linear model

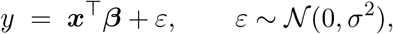

with feature vectors ***x*** ~ *N* (**0, Σ**) ∈ ℝ^*p*^, where **Σ** is a fixed population covariance matrix.

##### Population covariance

The covariance **Σ** is specified via its eigen-decomposition **Σ** = ***Q***Λ***Q***^⊤^, where ***Q*** ∈ ℝ^*p×p*^ is orthogonal and Λ = diag(*λ*_1_, …, *λ*_*p*_) contains the population eigenvalues. In different experiments we consider either spiked spectra or power-law spectra, for example *λ*_*j*_ ∝ *j*^−*α*^, with a global scaling chosen so that the average marginal variance has a prescribed value.

##### Signal vector, signal variance and noise variance

The ground-truth regression coefficient ***β*** ∈ ℝ^*p*^ is constructed to lie in the span of the top *k* eigenvectors of **Σ**. Let *Q*_*k*_ ∈ ℝ^*p×k*^ denote the first *k* columns of *Q* and let ***θ*** ∈ ℝ^*k*^ encode the signal in this principal subspace. We set ***β***_0_ = *Q*_*k*_***θ***, and then rescale to obtain the final coefficient vector 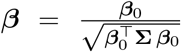. By construction this enforces ***β***^⊤^**Σ *β*** = 1, so the signal variance is normalized to one across all simulations in this setting. This normalization makes comparisons across different covariance structures and noise levels more interpretable, because changes in prediction error can be directly attributed to geometric and estimation effects rather than to a trivial rescaling of the signal strength. For different signal-to-noise ratio, we then set *σ*^2^ to different values accordingly.

##### Sampling training and test data

Given (**Σ, *β***), the training design matrix *X*_train_ ∈ ℝ^*n×p*^ has independent rows 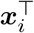 drawn from *N* (0, **Σ**), and the corresponding responses are

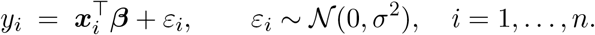

New test points ***x***_0_ and ***x***_new_ are always sampled independently from the same feature distribution *N* (0, **Σ**) and labeled via the same linear model.

#### S4.2 PCR estimator and effective projection matrix

Let *X*_train_ ∈ ℝ^*n×p*^ be the training design matrix with rows 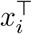 and assume the data are centered so that no intercept is needed. Define the empirical covariance

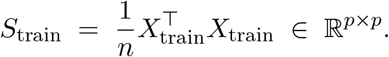

Let *V*_*k*_(*X*_train_) ∈ ℝ^*p×k*^ denote the matrix whose columns are the top *k* eigenvectors of *S*_train_, ordered by decreasing eigenvalue; when the dependence on *X*_train_ is clear we write *V*_*k*_ for brevity.

The *n × k* matrix of principal component scores is

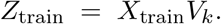

PCR performs ordinary least squares (OLS) of *y*_train_ on *Z*_train_. Let

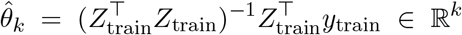

be the OLS coefficients in PC space, and define the corresponding coefficient vector in the original *p*-dimensional space by

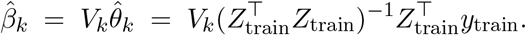

Using *Z*_train_ = *X*_train_*V*_*k*_ and 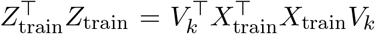, we can write 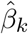 as a linear transformation of *y*_train_ in the original feature space:

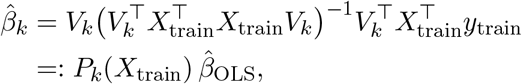

where 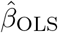 is the (hypothetical) OLS estimator on the full *p*-dimensional design, and the matrix

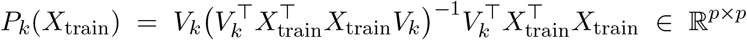

acts as an *effective projection* of *β* induced by PCR. This is the projection matrix that appears in the bias term and is more accurate than simply using the orthogonal projector 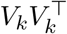.

Given a new feature vector *x*_0_ ∈ ℝ^*p*^, the PCR prediction is

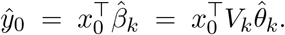

We analyze the mean squared prediction error 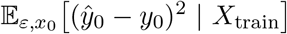 and decompose it into squared bias and variance terms, following the conditional-on-*X*_train_ approach of Green & Romanov (2025).

#### S4.3 Bias of PCR at a fixed test point

Recall that the responses follow 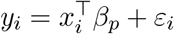 with independent *ε*_*i*_ ~ *N*(0, *σ*^2^). Here *β*_*p*_ and Σ_*p*_ (below) denote the population coefficient vector and covariance for this *p*-dimensional feature space. Conditional on *X*_train_ and a fixed test point *x*_0_, we have

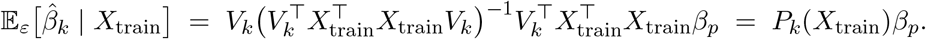

Therefore

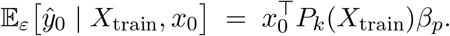

The conditional bias at *x*_0_ is

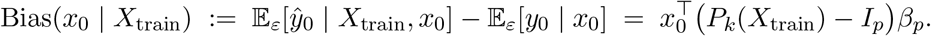

The squared bias at *x*_0_ is then

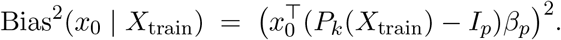

Averaging over a random test point *x*_0_ ~ *N*(0, **Σ**_*p*_), independent of *X*_train_, yields the integrated squared bias

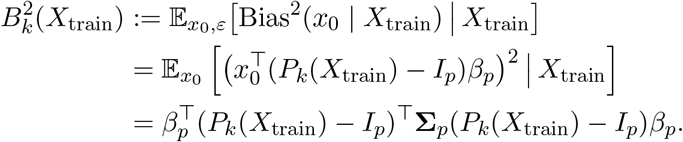

This is the quantity we estimate empirically in our simulations by averaging over many draws of *x*_new_ for each fixed training set *X*_train_.

#### S4.4 Variance of PCR at a fixed test point

We now compute the prediction variance due to the label noise *ε*. Write the PCR estimator as

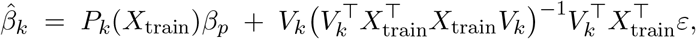

so that conditional on *X*_train_ we have

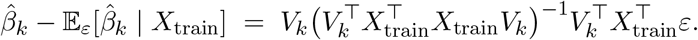

Since *ε* ~ *N* (0, *σ*^2^*I*_*n*_) independent of *X*_train_, the conditional covariance of 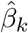 is

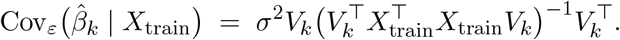

For a fixed test point *x*_0_ we obtain the conditional prediction variance

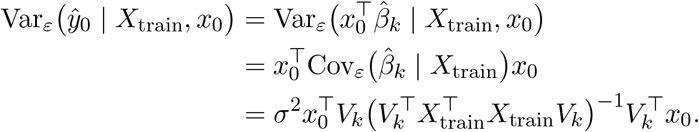

Averaging over a random test point *x*_0_ with covariance **Σ**_*p*_ gives the integrated variance term

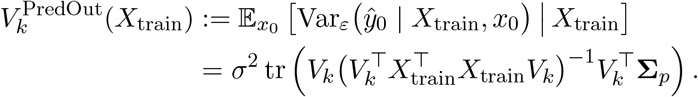

This is exactly the form used to interpret the out-of-sample variance in the main text, with *V*_*k*_ differing between the non-leaky and leaky settings.

#### S4.5 In-sample prediction variance

For completeness, we briefly record the in-sample prediction variance, which is simpler and depends only on the number of retained components *k*. Let *H*_*k*_ be the *n × n* PCR hat matrix,

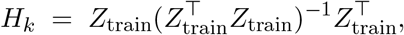

so that *ŷ*_train_ = *H*_*k*_*y*_train_. Conditional on *X*_train_, the in-sample prediction variance averaged over the *n* training points is

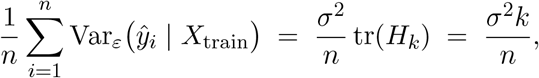

since *H*_*k*_ is an idempotent matrix of rank *k*. This recovers the well-known result that the in-sample variance term for PCR depends only on the number of PCs used, not on *p* or on **Σ**_*p*_.

#### S4.6 Out-of-sample variance

We now rewrite the variance expression from the previous subsection in a notation that matches the out-of-sample prediction risk decomposition of Green & Romanov (2025).

Recall that for a new test point ***x***_0_ ~ *N* (0, Σ_*p*_), independent of (*X*_train_, *y*_train_), the PCR prediction is 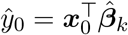. Conditional on *X*_train_ and ***x***_0_ we showed that

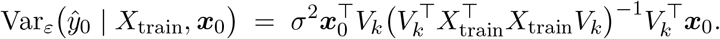

Averaging over the test covariate ***x***_0_ ~ *N* (0, Σ_*p*_) yields the *out-of-sample variance term*

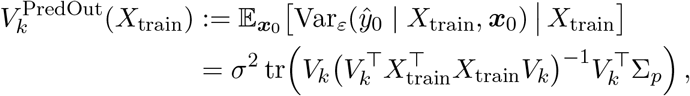

which is exactly the variance component of the conditional out-of-sample prediction risk considered in Green & Romanov (2025), specialized to our finite sample setting.

To make contact with the notation in Green & Romanov (2025), it is convenient to introduce the rescaled sample covariance 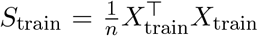 and write 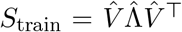 with eigenvalues 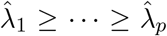 and eigenvectors 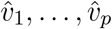. Then 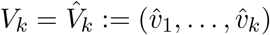 and 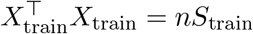, so that

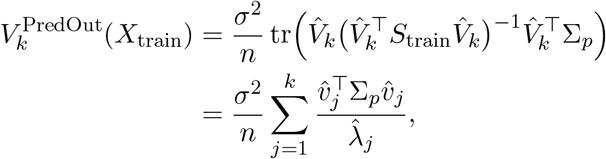

which matches the empirical out-of-sample variance term 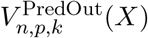 in Green & Romanov (2025). In the isotropic case Σ_*p*_ = *I*_*p*_, this simplifies to

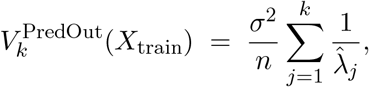

so the out-of-sample variance is fully determined by the inverse empirical eigenvalues associated with the retained principal components.

#### S4.7 Comparison of in-sample and out-of-sample variance

We now summarize the relationship between the in-sample and out-of-sample variance terms derived above.

##### Finite-sample expressions

The average in-sample prediction variance of PCR, conditional on *X*_train_, is

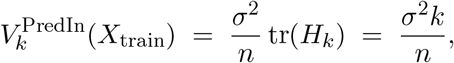

since the PCR hat matrix *H*_*k*_ has rank *k*. In contrast, the out-of-sample variance term is

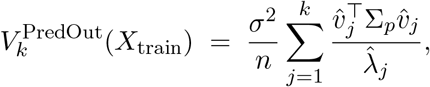

which depends on the interaction between the empirical PC directions 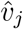, their empirical eigenvalues 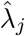, and the population covariance Σ_*p*_. Thus 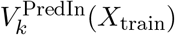 depends only on *k* and *n*, whereas 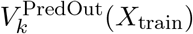 encodes detailed spectral information about both the sample and population covariance.

##### Special case: isotropic features

When Σ_*p*_ = *I*_*p*_, we have

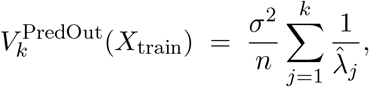

so 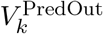 can be compared directly to 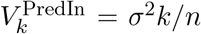 via the empirical spectrum 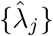. If the top *k* sample eigenvalues are overestimated relative to their population counterparts (as is typical for leading components in high-dimensions), then 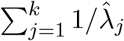 tends to be smaller than *k*, and 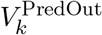 is often below 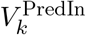, although no strict ordering holds uniformly across all covariance structures.

##### High-dimensional asymptotics

In the proportional asymptotic regime of Green & Romanov (2025), where *p/n* → *γ* and *k/p* → *α*, the in-sample variance remains *σ*^2^*k/n* while the out-of-sample variance converges almost surely to a spectral functional 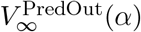 that depends on the limiting eigenvalue distribution of Σ_*p*_ and on the eigenvector overlaps between the signal and the population PCs. In particular, 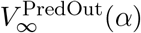 can be strictly above or below 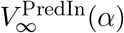 depending on the covariance model, illustrating that in-sample variance is not a universal upper bound for out-of-sample variance in high dimensions.

#### S4.8 Leaky vs. non-leaky PCR variance term

The derivations above hold for any PCA basis *V*_*k*_ and therefore apply to (i) the non-leaky basis *V*_*k*_(*X*_train_) computed from the training data only and (ii) the leaky basis *V*_*k*_(*X*_aug_) computed from the augmented design 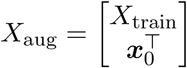 that includes the test point. For a fixed training set, we compare three variance terms:

- In-sample variance 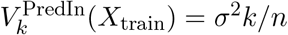, which depends only on *k* and *n*.
- Out-of-sample (non-leaky) variance 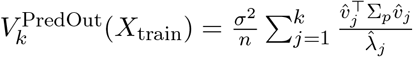, with *V*_*k*_ computed on *X*_train_.
- Leaky CV variance, obtained when the PCA basis is computed on *X*_aug_ and the prediction is evaluated at the same test point ***x***_0_ used in the basis:

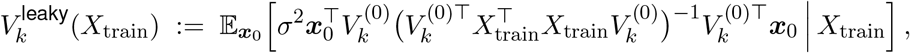

where 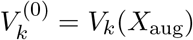.

In the leaky setting, the PCA basis 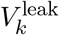 is computed from an augmented design matrix *X*_aug_, which includes the test point ***x***_0_. The regression is still fit only on the *n* training responses, but the PCA basis has implicitly “seen” the test covariate. We compare the resulting out-of-sample variance term at ***x***_0_ to the in-sample prediction variance.

The key observation is that the Gram matrix of the augmented design can be written as

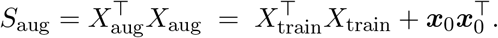

Let 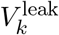 denote the top *k* eigenvectors of *S*_aug_, with eigenvalues *λ*_1_ ≥ · · · ≥ *λ*_*k*_, and define the leverage of ***x***_0_ in this PC subspace as

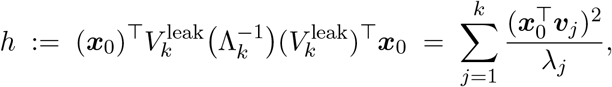

where Λ_*k*_ = diag(*λ*_1_, …, *λ*_*k*_) and ***v***_*j*_ are the corresponding eigenvectors. This quantity measures how much ***x***_0_’s position in predictor space influences its own fitted value—the standard definition of statistical leverage in regression.

Using the Sherman–Morrison formula to invert the rank-1 perturbation 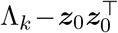 (where 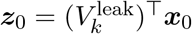 is the score of ***x***_0_ in the leaky PC basis), we obtain an exact expression for the leaky out-of-sample variance:

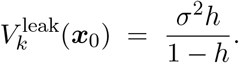

Comparing this to the in-sample variance 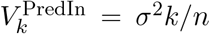, we see that the leaky variance is larger whenever

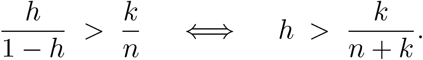

Since the average leverage across the *n* + 1 augmented points is 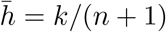, and since *k/*(*n* + 1) *> k/*(*n* + *k*) for any *k >* 1, the test point ***x***_0_ drawn from the same distribution as the training data will typically have leverage above the threshold, and thus exhibit a leaky variance strictly larger than the in-sample term. This inflated variance is a direct consequence of the data leakage: by including ***x***_0_ in the unsupervised PCA step, the PCA basis becomes slightly “aligned” to ***x***_0_, increasing its effective dimensionality in the retained PC subspace.

##### Power-law spectrum

Under power-law eigenspectra for Σ_*p*_, our simulations showed a strict ordering:

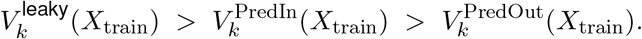

Intuitively, power-law spectra concentrate variance in leading components; the leaky basis over-aligns these components with ***x***_0_, further inflating its leverage and hence the variance term.

##### Generalization variance and the non-leaky term

The variance component of the true generalization error for the leaky model (same leaky basis, but evaluated at an independent ***x***_new_) closely matched the non-leaky out-of-sample variance 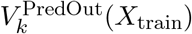 in our experiments. This is expected because adding a single point to form *X*_aug_ perturbs the basis by *O*(1*/n*), whereas both generalization terms integrate over independent test covariates.

##### No universal ordering

Although in the power-law case we found 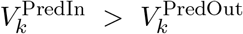, in-sample variance is not a universal upper bound for out-of-sample variance. Depending on the covariance structure (e.g., spiked or isotropic models) and finite-sample geometry, 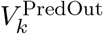 can be above or below 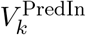.

#### S4.9 Varying number of features via a master model

So far in this section, the primary simulation setting assumes a fixed feature dimension *p* and focuses on how PCR behaves as the number of training samples *n* and the PCA dimensionality *k* are varied. We next extend this setup to the case where *p* increases while *n* is held fixed, and examine how PCR’s bias and variance terms change with dimensionality. To do this in a way that preserves a coherent underlying signal, we adopt a “master model” construction and obtain lower-dimensional problems by subsampling coordinates, rather than regenerating a new ***β*** for each value of *p*.

##### Master covariance and signal

We fix a large dimension *P*_max_ and construct a master covariance 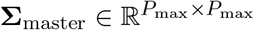 with a power-law eigenspectrum, together with a master coefficient vector 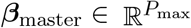 that lies in the span of the top *k* master eigenvectors. Exactly as above, ***β***_master_ is scaled so that 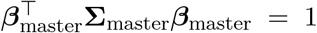, ensuring that the signal variance of the underlying master model is one.

##### Subsampling to obtain different feature dimensions

For any target feature dimension *p* ≤ *P*_max_ we form the *p × p* principal submatrix 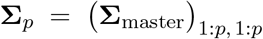 and define the corresponding regression vector by truncation, 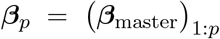 and then scaled such that 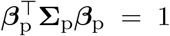. Training and test samples in dimension *p* are then drawn as ***x***_*i*_ ~ *N* (0, **Σ**_*p*_) with 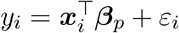 as before.

##### Subsampling vs regenerating *β*_*p*_ for each subspace

A common approach in high-dimensional simulation studies is to re-draw a new ***β***_*p*_ independently for each ambient dimension *p*, often again supported on the top eigenvectors of **Σ**_*p*_. While convenient, this changes the true signal subspace each time *p* changes, so differences in performance conflate genuine dimensional effects with changes in the underlying regression function. Our subsampling construction keeps a single master signal ***β***_master_ fixed and views smaller-*p* problems as coordinate restrictions of the same latent model. This has two advantages: (i) it aligns more closely with the scientific interpretation that increasing *p* reveals additional measured features of the same underlying system, and (ii) it allows us to attribute changes in PCR bias and variance directly to spectral and estimation effects induced by increasing dimension, rather than to changes in the ground-truth ***β***.

#### S4.10 Multiple leaked points

So far we have considered the behavior of PCR MSE when there was one leaked point. Here we extend this setup by allowing multiple leaked points, and derived a bound between the leaky estimated variance term on the leaked points, and in-sample variance.

##### Setup: augmented design and PC basis

When multiple points are included in the augmented design matrix, let *X*_leak_ ∈ ℝ^*m×p*^ denote the *m* leaked points. The augmented design is

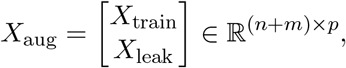

and the corresponding Gram matrix is 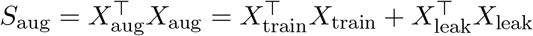.

Let 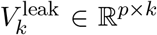 denote the matrix of top *k* eigenvectors of *S*_aug_, with corresponding eigenvalues *λ*_1_ ≥ · · · ≥ *λ*_*k*_ *>* 0 collected in Λ_*k*_ = diag(*λ*_1_, …, *λ*_*k*_). The eigendecomposition gives 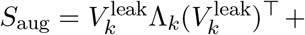 (lower rank terms).

##### Score matrices and the leaky Gram matrix

Define the *n × k* and *m × k* matrices of PC scores:

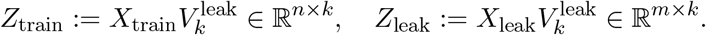

In the leaky PC basis, the Gram matrices of the training and leaked data satisfy 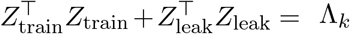. Therefore, the Gram matrix of the training data alone in this basis is 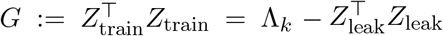. This is a rank-*m* perturbation of the diagonal matrix Λ_*k*_.

##### PCR prediction at leaked points

The PCR estimator fits OLS on *Z*_train_ against training labels **y**_train_, obtaining coefficients 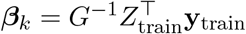. The predicted labels at the *m* leaked points are

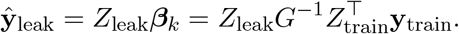

Since **y**_train_ has covariance *σ*^2^*I*_*n*_ (independent noise), the covariance of the predictions is

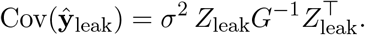

##### Block matrix inversion

To compute *G*^−1^, we write *G* as

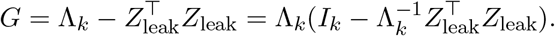

then apply the Woodbury matrix identity. Thus,

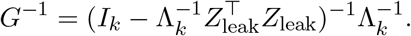

Define the *m × m* matrix

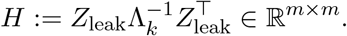

This is the block leverage matrix of the leaked points in the leaky PC basis. Note that *H* is symmetric and positive semi-definite. By the Woodbury identity applied to (*I*_*k*_ − *H*^*′*^)^−1^ where 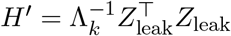,

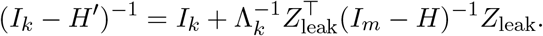

Therefore

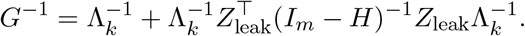

Now compute 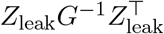.

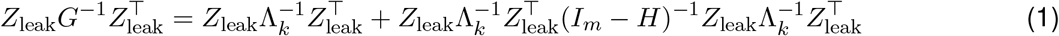

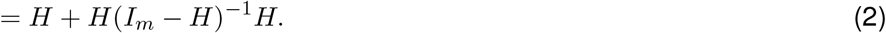

Since *H* is symmetric, we use the identity *H* + *H*(*I*_*m*_ − *H*)^−1^*H* = *H*(*I*_*m*_ − *H*)^−1^. Therefore

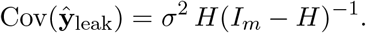

##### Variance at individual leaked points

The diagonal entries of Cov(***ŷ***_leak_) give the variance at each leaked point:

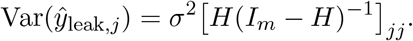

The sum of all variances is

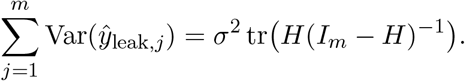

Let *η*_1_, …, *η*_*m*_ denote the eigenvalues of *H*. These lie in (0, 1) because (i) *H* is a leverage matrix (non-negative), and (ii) the invertibility of *G* ensures that *η*_max_ *<* 1. Then

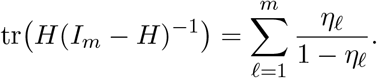

The average variance per leaked point is

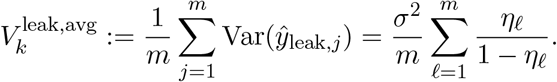

##### Comparison to in-sample variance

Recall from Section S4.5 that the average in-sample PCR variance per training point is

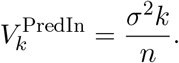

To compare 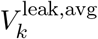 to 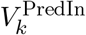, we use Jensen’s inequality. The function *f*(*η*) = *η/*(1 − *η*) is strictly convex on (0,1), therefore

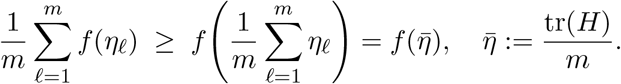

This gives

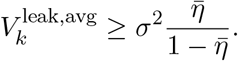

##### Expected leverage budget

The total leverage in any regression onto *k* dimensions is exactly *k*. Specifically, summing over all *n* + *m* rows (training and leaked),

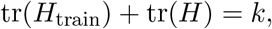

where 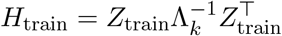 is the leverage matrix of training points.

For a random sample of *m* i.i.d. points from the same distribution as the training data, each point has expected leverage *k/*(*n* + *m*), so on average

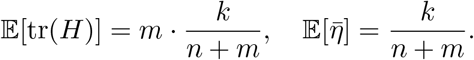

Therefore

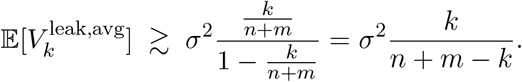

##### The threshold condition: m < k

Comparing the expected average leaky variance to the in-sample variance:

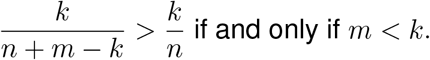

##### Conclusion

The average leaky variance per leaked point exceeds the in-sample variance per training point whenever m<k.

This result generalizes the single-point case: for m=1, the condition becomes k>1, which recovers the finding in Section S4.8 that a single leaked point typically exhibits inflated variance (provided k>1). For multiple leaked points, the threshold naturally scales with m: more leaked points “dilute” their collective leverage, reducing the average variance inflation unless their number is small relative to the number of PC components.

### S5 Temporal-covariance leakage correction: scalar and full procedures

#### Notation

| Symbol | Meaning |
| --- | --- |
| $k$ | Class index ($k \in \{1, 2\}$) |
| $K$ | Number of test samples |
| $n$ | Number of training time points per class |
| $\mu$ | Class mean ($\mu_1 = \mu, \mu_2 = -\mu$) |
| $A$ | Within-training covariance block |
| $C$ | Train–test cross-covariance block |
| $c_{\text{eff}}$ | Class-averaged covariance reduction |
| $\Phi(\cdot)$ | Cumulative probability distribution for the standard Gaussian |
| $P_{\text{gen}}, P_{\text{train}}, P_{\text{test}}$ | Generalization / training / leaky test error |

This section provides additional details on the two correction variants applied in Fig. 6h. Both aim to recover the generalization error of a classifier trained on a temporally correlated signal from the (downward-biased) leaky test error produced by a train–test split that does not respect trial boundaries.

#### S5.1 Problem setup and parameter estimation

To formally demonstrate how temporal dependencies alter performance metrics, we consider a simple binary discrimination task using a univariate linear discriminant analysis (*p* = 1, 2 classes: *k* ∈ *{*1, 2*}*). In this setting, the neuron response fluctuates around two scalar class means *µ*_1_ and *µ*_2_, with a temporal covariance structure. Specifically, for a class with *n*_full_ time points, the response vector 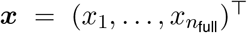 follows a multivariate Gaussian distribution ***x*** ~ *N* (*µ*_*k*_**1**, *A*_full_). According to a specified train-test split, we can partition the covariance matrix *A*_full_ into four blocks:

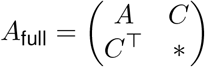

where *A* is the covariance within the training set, and *C* is the cross-covariance between the training and test sets.

The train set is ***X***_**1**_ = *µ*_*k*_**1** + ***ϵ***_**1**_, where ***ϵ***_**1**_ ~ *N*(0, *A*). The test set is ***X***_**2**_ = *µ*_*k*_**1** + ***ϵ***_**2**_, where the cross-covariance between train and test sets is Cov(***ϵ***_**1**_, ***ϵ***_**2**_) = *C*. For simplicity, we assume:

- The two classes are balanced, and the covariance structure is identical across classes, so we drop the class index *k*.
- The noise distribution is stationary across the train and the test sets, so the noise in the test set has marginal variance 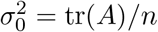.
- The class means are symmetric around zero, i.e., *µ*_1_ = *µ, µ*_2_ = −*µ*.

The LDA classifier is determined by the naive Bayes decision rule:

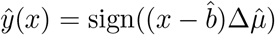

where 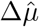 is the mean difference between the two classes and 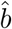 is the decision boundary.

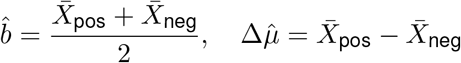

Note that samples are correlated within the train set. Defining the variance of the estimated decision boundary as 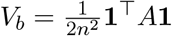, we have

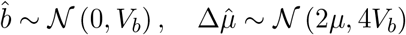

#### S5.2 Generalization error and training error

Applying the model to an independent test sample *x* = *µ* +*ϵ* where 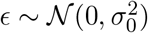, the generalization error is given by the Gaussian cumulative distribution function Φ. Since *x* and 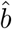 are independent,

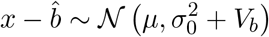

Conditioning on the estimated discriminant direction being correct 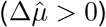, the true generalization error is given by:

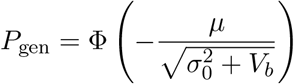

For the in-sample prediction error, 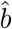 and the training samples are dependent. For the positive training samples *x*_*i*_ = *µ* + *ϵ*_*i*_, *i* = 1, …, *n*, the error is given by 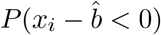. The variance of 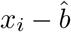 is:

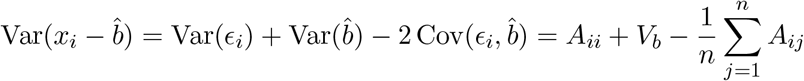

So the training error rate is given by:

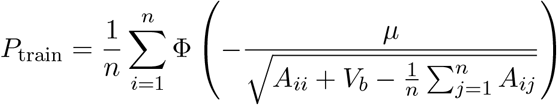

Note that if *A* is positive definite then *V*_*b*_ *>* 0, so the average training prediction variance 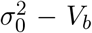 is strictly below the generalization variance 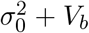, and the training error is correspondingly smaller than the generalization error.

#### S5.3 Test error and leakage bounds

A simple expansion reveals the impact of data leakage on the generalization error estimation from the test set. Considering a non-zero cross-covariance *C*, the finite *K*-sample test error is:

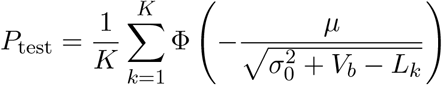

where 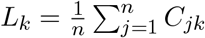 is the covariance reduction for the *k*-th test sample.

If the sum of all entries in *C* normalized by the test size *K* equals some constant *c*, we can bound the expected test error using Jensen’s inequality. Because the function 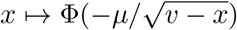 is convex when *v* − *x* ≤ *µ*^2^*/*3, applying this convexity condition (which corresponds to a sufficiently strong signal-to-noise ratio, SNR) yields:

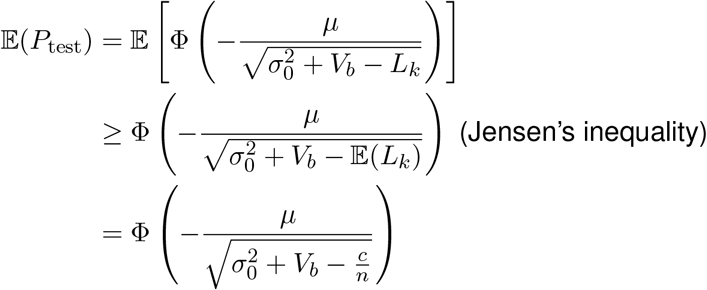

The introduction of Jensen’s inequality removes the variation in the averaging *L*_*k*_, which fluctuates around *c/n*. If *L*_*k*_ concentrates around *c/n* with high probability, under the Lipschitz condition, we can get a high probability bound of E(*P*_test_). Assuming that for some *ϵ, δ >* 0,

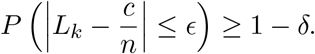

Moreover, suppose that the denominator is bounded away from zero, i.e., for all 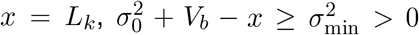. Then the map 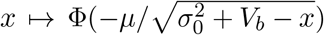 is Lipschitz with constant 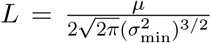. Therefore, with probability at least 1 − *δ*,

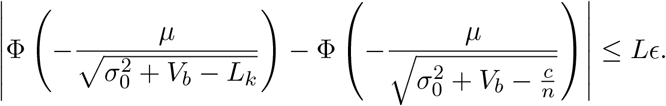

Hence, if the per-sample leakage *L*_*k*_ concentrates around *c/n*, the test error also concentrates around its deterministic approximation.

#### S5.4 Extension to interleaved class labels

When positive and negative trials are interleaved on a single time axis, the two class-conditional noise vectors share a common temporal process, and the single cross-covariance block *C* no longer suffices. Let *P*_tr_, *N*_tr_, *P*_te_, *N*_te_denote the time indices of positive training, negative training, positive test, and negative test samples, each of size *n, n, K, K* respectively. Define the class-resolved train–test cross-covariance blocks

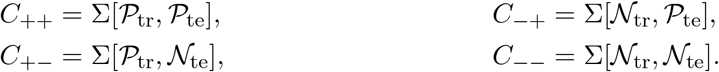

Because the decision boundary 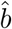 is estimated from both classes, each test sample’s effective leakage involves covariance with all training samples. The per-sample covariance reductions are

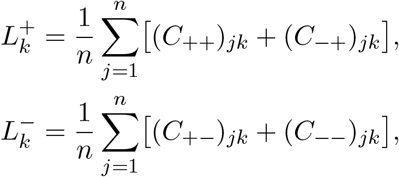

and the class-averaged scalar leakage is

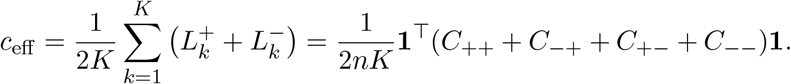

Positive *c*_eff_reduces the apparent test variance and biases the test error downward.

#### S5.5 Scalar correction

The scalar correction treats all held-out samples as having the same leakage, equal to *c*_eff_. Given an observed leaky test error 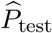, we first convert it to an apparent prediction variance,

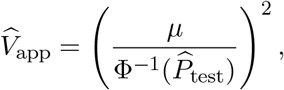

and then add the scalar leakage back:

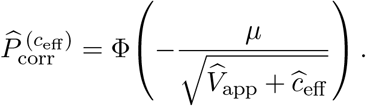

This closed-form correction shifts the apparent variance upward by the amount that the train–test covariance artificially reduced it. The advantage of the scalar correction is its simplicity: only a single number, 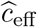, needs to be estimated from the temporal covariance.

#### S5.6 Full per-sample correction

When the per-sample leakage values 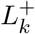 and 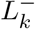 are heterogeneous (as occurs with interleaved or irregularly ordered class labels), the scalar approximation can be imprecise because the error function Φ is nonlinear. The full correction retains the complete leakage vector and solves for the corrected generalization variance *V* ^∗^ that satisfies

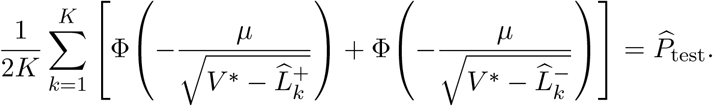

Because the left-hand side is monotonically increasing in *V* ^∗^ (for 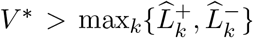), we solve this equation numerically via Brent’s root-finding method. The corrected generalization error is then

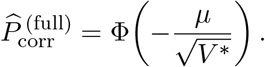

By accounting for per-sample variation in the train–test covariance, the full correction can be more accurate than the scalar correction when the leakage landscape is uneven across held-out samples.

#### S5.7 Covariance estimation

Both the above corrections require an estimate of the temporal covariance matrix Σ. We assume a stationary exponential model, Σ_*ij*_= *σ*^2^*α*^|*i*−*j*|^ with 0 *< α <* 1, and estimate the parameters *σ*^2^ and *α* from residual activity (after subtracting class means) using a semivariogram-based fit. The empirical half-squared-increment curve,

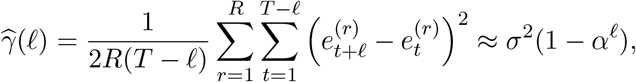

is fitted by weighted least squares over lags *ℓ* = 1, …, *L*, with weights *w*_*ℓ*_= *T* − *ℓ*. This estimator is more robust than the raw autocovariance for short, highly autocorrelated traces because the half-squared increment does not require explicit mean removal (which can bias long-lag autocovariances downward). The variance estimate is mildly regularized by blending the semivariogram-derived variance with the empirical residual variance (see Methods). Once 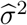 and 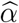 are obtained, the estimated covariance matrix is indexed by the original train and test sample positions to compute 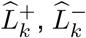, and 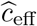.

#### S5.8 Summary

Temporal autocorrelations intrinsically violate the independence assumptions of standard cross-validation, producing a predictable, downward-biased estimate of generalization error. As we have shown, this bias is driven by the cross-covariance between the training and test sets. By modeling this temporal covariance structure, and applying either the simple scalar correction or the more precise full per-sample correction, we can analytically reverse the artificial variance reduction. These correction procedures thus offer a practical tool for reporting unbiased decoding performance in continuous neural recordings without needing to discard large segments of data to enforce strict independence.

---

**Figure S1:**
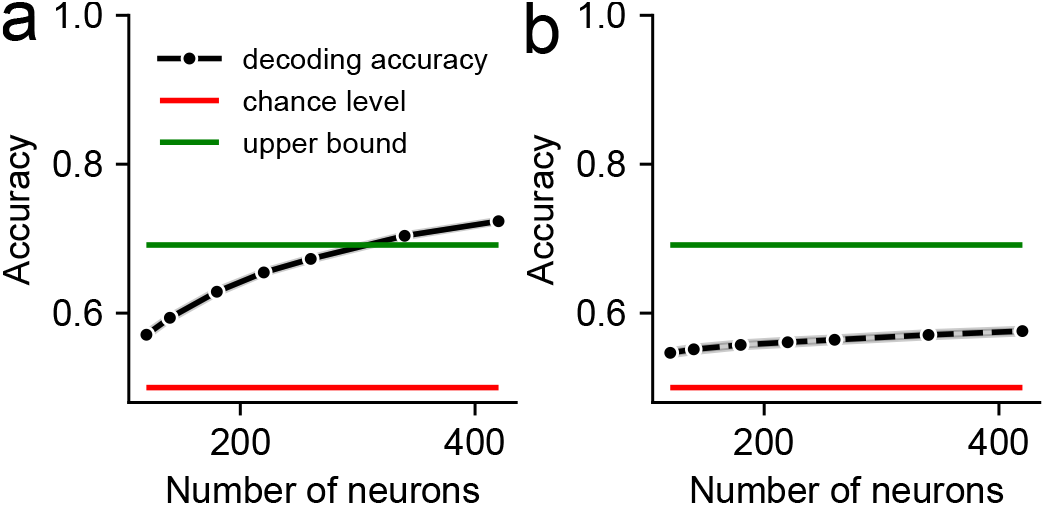
Neurons without the Gaussian parametric tuning curves assumption. **(a)**: Similar to Fig. 2c. **(b)**: Similar to Fig. 2d.

**Figure S2:**
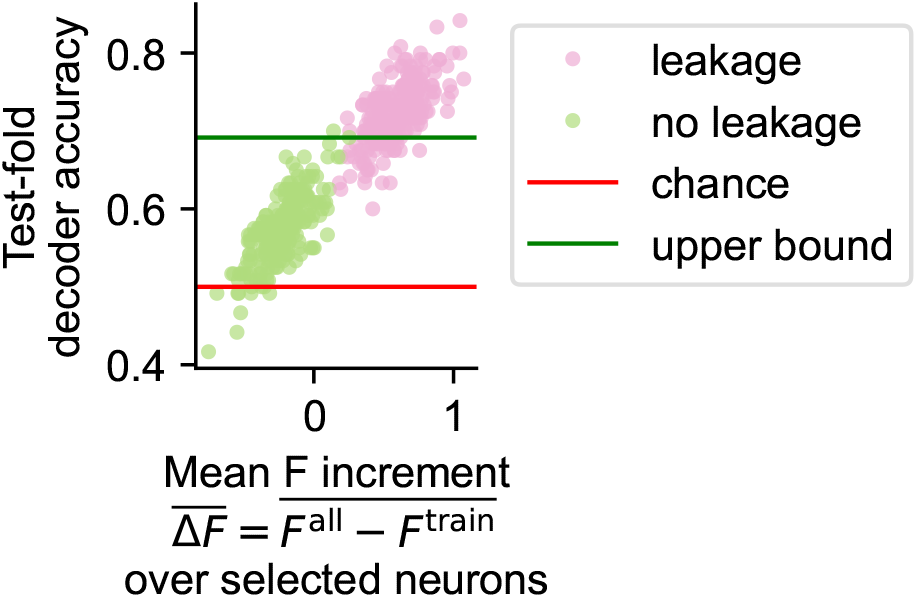
Inflated decoding accuracy tracks the leaked F-statistic increment of the selected neurons. Neural activity was simulated as in Fig. 2d using a Gaussian tuning-curve generative model (N=420 neurons, M=300 trials per class, 5-fold stratified cross-validation, k=100 selected neurons, 50 independent simulations). For each cross-validation fold, neurons were ranked two ways: (i) leaky selection (pink), top-k neurons by the F-statistic *F* ^all^ computed on all trials, and (ii) clean selection (green), top-k neurons by the F-statistic *F*^train^ computed on the training trials only. For each selection, we computed the per-neuron *leaked F increment* 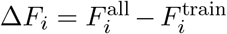 and averaged it over the *k* selected neurons (*x*-axis). This quantity measures how much including the held-out trials in the ranking pushed a selected neuron’s score above its train-only value. The *y*-axis shows the LDA test-fold decoding accuracy for that fold. Each dot is a single (simulation, fold) pair. Red line: chance accuracy (0.5); green line: theoretical upper bound for a linear decoder. Under leaky selection, 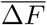 is positive and covaries with inflated accuracy, confirming that the leaked component of the ranking drives the accuracy inflation. Under non-leaky selection, 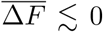 (independent test trials dilute the train-optimized score), and accuracy is between the upper bound and chance level.

**Figure S3:**
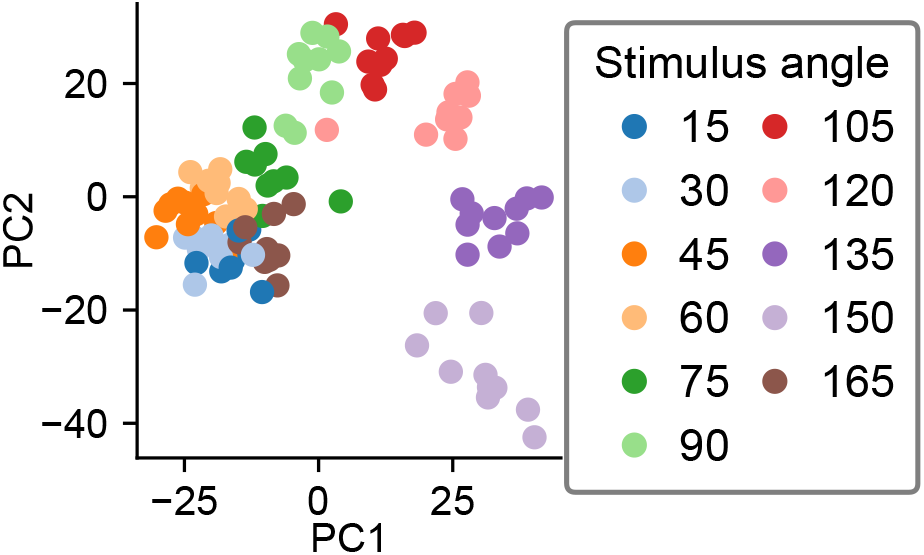
Similar to Fig. 3d, PC embedding using trial-averaged PCA of the contralateral activity. Each dot is one trial.

**Figure S4:**
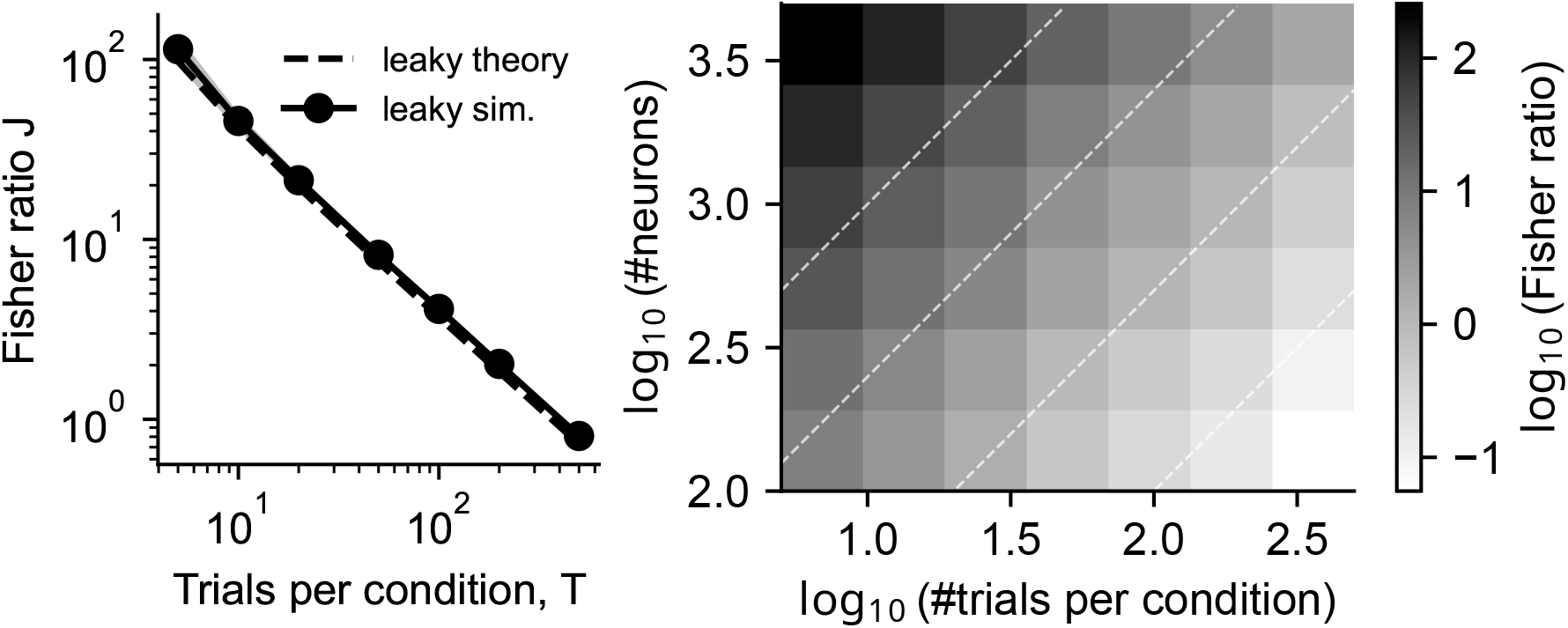
Apparent cluster separability under global trial-averaged PCA scales with *d/T*. Random neural activity (i.i.d. *N* (0, 1), no true signal) was generated with *C* = 11 conditions. Trial-averaged PCA was fit on the full dataset and all *C* × *T* trials were projected onto the top *k* = 2 PCs. Cluster separability was quantified by the multiclass Fisher discriminant ratio (*J*), higher *J* indicates better apparent separability. Results are shown as mean ± s.d. over 100 random seeds. **(a)**: *J* as a function of trials per condition *T* at fixed *d* = 2000. The dashed line shows the analytical prediction derived in Section S2. **(b)**: Heatmap of log_10_*J* over a grid of (*d, T*) values (mean over 100 seeds). Dashed white lines mark constant *d/T* ratios; the iso-*J* contours align with these diagonals, showing that the apparent clustering is governed by *d/T* rather than by *d* or *T* alone.

**Figure S5:**
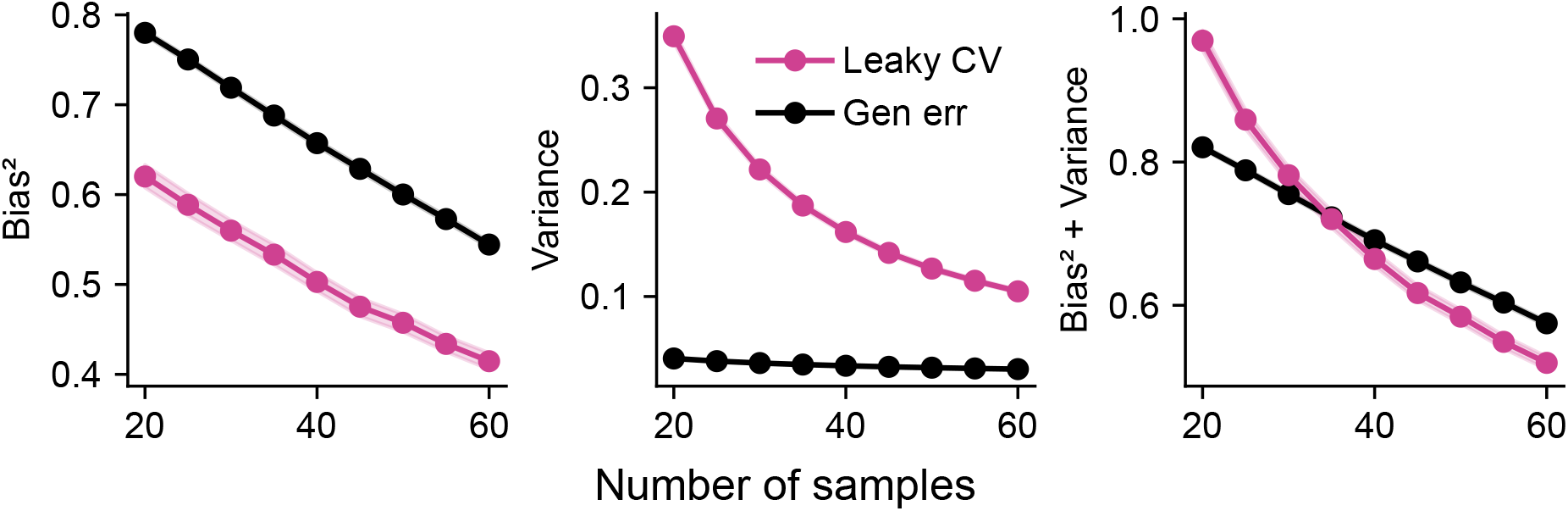
Similar to Fig. 5e but with spiked covariance for the signal variance.

**Figure S6:**
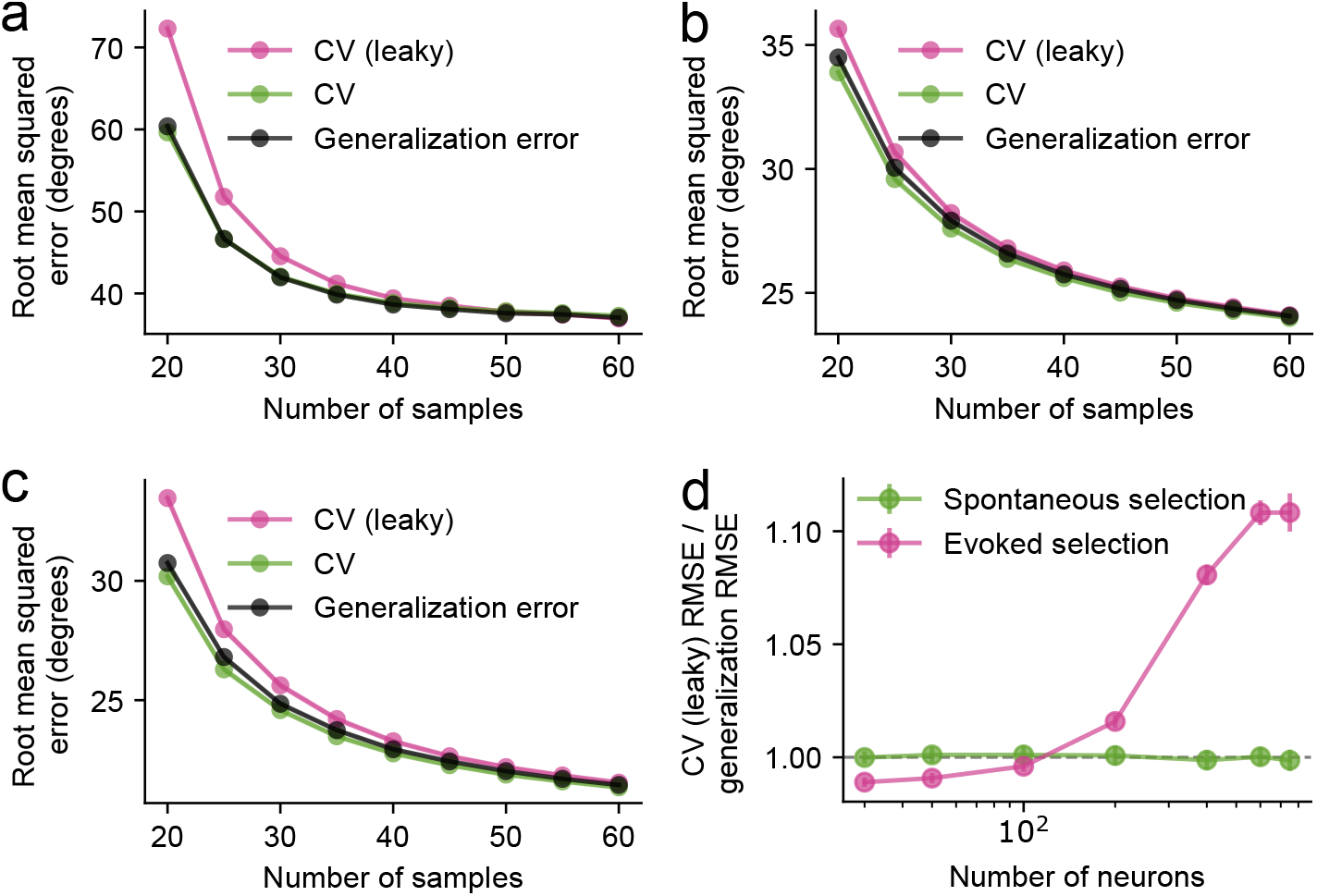
Unsupervised feature selection shows biased results compared to true generalization error. **(a)**: Similar to Fig. 4c. Gaussian tuning-curve responses with neurons ranked by mean evoked firing rate before leave-one-out cross-validation. **(b)**: Similar to Fig. 4c with semi-parametric Gamma-Poisson response model with neurons ranked by mean evoked firing rate. Cross-validated error from the leaky model was higher than its generalization error. The non-leaky model closely tracked the generalization error, despite having one less sample used in the feature selection step. **(c)**: Similar to (b), ranked by evoked spike-count variance. In (a–c), globally selecting neurons before cross-validation makes the leaky CV RMSE diverge from the true generalization RMSE of the same leaky model, whereas non-leaky CV closely tracks generalization error. **(d)**: Similar to Fig. 4d with selection of the top 10% of neurons by mean firing rate.

**Figure S7:**
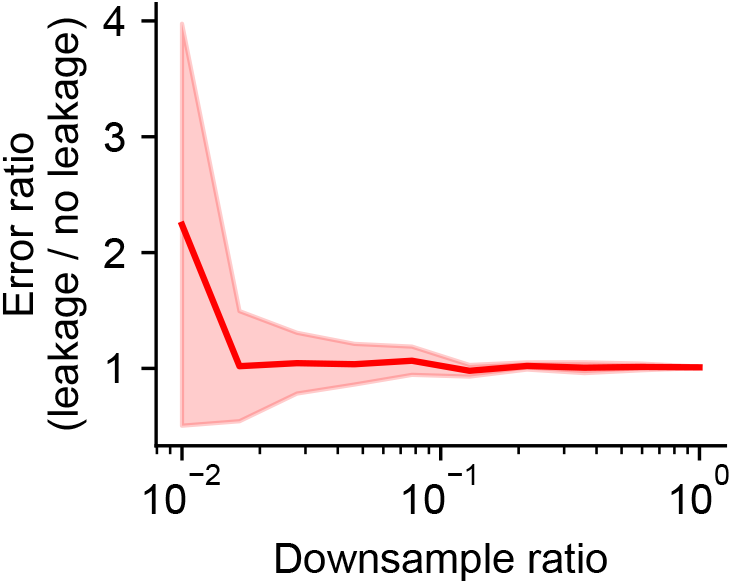
Downsampling real neural recordings increases the error mismatch from global z-scoring. We repeated the z-scoring-only decoding analysis from Stringer et al. (2021) after downsampling each session to different fractions of its original stimulus presentations. The plotted quantity is the ratio between the median decoding error from global, leaky z-scoring and the median decoding error from z-scoring refit within each cross-validation fold. Values above one indicate higher error under global z-scoring. The mismatch increased as fewer samples were retained, showing that leakage effects become more pronounced in smaller-sample regimes.

## References

Abu-Mostafa, Y. S., Magdon-Ismail, M., & Lin, H.-T. (2012). Learning from data, vol. 4. (AMLBook New York).

Ahrens, M. B., Orger, M. B., Robson, D. N., Li, J. M., & Keller, P. J. (2013). Whole-brain functional imaging at cellular resolution using light-sheet microscopy. Nature Methods, 10, 413–420.

Ambroise, C. & McLachlan, G. J. (2002). Selection bias in gene extraction on the basis of microarray gene-expression data. Proceedings of the National Academy of Sciences, 99, 6562–6566.

Anumanchipalli, G. K., Chartier, J., & Chang, E. F. (2019). Speech synthesis from neural decoding of spoken sentences. Nature, 568, 493–498.

Azabou, M., Arora, V., Ganesh, V., Mao, X., Nachimuthu, S. B., Mendelson, M. J., Richards, B. A., Perich, M. G., Lajoie, G., & Dyer, E. L. (2023). A unified, scalable framework for neural population decoding. In Thirty-seventh Conference on Neural Information Processing Systems.

Azabou, M., Pan, K. X., Arora, V., Knight, I. J., Dyer, E. L., & Richards, B. A. (2025). Multi-session, multi-task neural decoding from distinct cell-types and brain regions. In The Thirteenth International Conference on Learning Representations.

Birbaumer, N. & Cohen, L. G. (2007). Brain-computer interfaces: communication and restoration of movement in paralysis. The Journal of Physiology, 579, 621–636.

Borst, A. & Theunissen, F. E. (1999). Information theory and neural coding. Nature Neuroscience, 2, 947–957.

Button, K. S. (2019). Double-dipping revisited. Nature Neuroscience, 22, 688–690.

Cawley, G. C. & Talbot, N. L. (2010). On over-fitting in model selection and subsequent selection bias in performance evaluation. J. Mach. Learn. Res., 11, 2079–2107.

Chen, Z., Qing, J., Xiang, T., Yue, W. L., & Zhou, J. H. (2023a). Seeing Beyond the Brain: Conditional Diffusion Model with Sparse Masked Modeling for Vision Decoding. In 2023 IEEE/CVF Conference on Computer Vision and Pattern Recognition (CVPR), pp. 22710–22720. (Los Alamitos, CA, USA: IEEE Computer Society).

Chen, Z., Qing, J., & Zhou, J. H. (2023b). Cinematic mindscapes: high-quality video reconstruction from brain activity. In Proceedings of the 37th International Conference on Neural Information Processing Systems, NIPS ‘23. (Red Hook, NY, USA: Curran Associates Inc.).

Cowley, B. R., Snyder, A. C., Acar, K., Williamson, R. C., Yu, B. M., & Smith, M. A. (2020). Slow drift of neural activity as a signature of impulsivity in macaque visual and prefrontal cortex. Neuron, 108, 551–567.e8.

Cunningham, J. P. & Yu, B. M. (2014). Dimensionality reduction for large-scale neural recordings. Nature Neuroscience, 17, 1500–1509.

Dayan, P. & Abbott, L. (2005). Theoretical Neuroscience. (MIT Press).

Degenhart, A. D., Bishop, W. E., Oby, E. R., Tyler-Kabara, E. C., Chase, S. M., Batista, A. P., & Yu, B. M. (2020). Stabilization of a brain–computer interface via the alignment of low-dimensional spaces of neural activity. Nature Biomedical Engineering, 4, 672–685.

Denker, M., Yegenoglu, A., & Grün, S. (2018). Collaborative HPC-enabled workflows on the HBP Collaboratory using the Elephant framework. Neuroinformatics 2018, Montreal (Canada), 9 Aug 2018-10 Aug 2018.

Ebrahimi, S., Lecoq, J., Rumyantsev, O., Tasci, T., Zhang, Y., Irimia, C., Li, J., Ganguli, S., & Schnitzer, M. J. (2022). Emergent reliability in sensory cortical coding and inter-area communication. Nature, 605, 713–721.

Ferri, F., Pudil, P., Hatef, M., & Kittler, J. (1994). Comparative study of techniques for large-scale feature selection. In Machine Intelligence and Pattern Recognition. (Elsevier), pp. 403–413.

Flesher, S. N., Downey, J. E., Weiss, J. M., Hughes, C. L., Herrera, A. J., Tyler-Kabara, E. C., Boninger, M. L., Collinger, J. L., & Gaunt, R. A. (2021). A brain-computer interface that evokes tactile sensations improves robotic arm control. Science, 372, 831–836.

Friedrich, J., Zhou, P., & Paninski, L. (2017). Fast online deconvolution of calcium imaging data. PLOS Computational Biology, 13, e1005423.

Gebhardt, C., Auer, T. O., Henriques, P. M., Rajan, G., Duroure, K., Bianco, I. H., & Del Bene, F. (2019). An interhemispheric neural circuit allowing binocular integration in the optic tectum. Nature Communications, 10, 5471.

Giovannucci, A., Friedrich, J., Gunn, P., Kalfon, J., Brown, B. L., Koay, S. A., Taxidis, J., Najafi, F., Gauthier, J. L., Zhou, P., Khakh, B. S., Tank, D. W., Chklovskii, D. B., & Pnevmatikakis, E. A. (2019). CaImAn an open source tool for scalable calcium imaging data analysis. eLife, 8, e38173.

Glaser, J. I., Benjamin, A. S., Chowdhury, R. H., Perich, M. G., Miller, L. E., & Kording, K. P. (2020). Machine learning for neural decoding. eneuro, 7, ENEURO.0506–19.2020.

Green, A. & Romanov, E. (2025). The high-dimensional asymptotics of principal component regression. The Annals of Statistics, 53, 1697–1727.

Grosmark, A. D. & Buzsáki, G. (2016). Diversity in neural firing dynamics supports both rigid and learned hippocampal sequences. Science, 351, 1440–1443.

Hadidi, N., Feghhi, E., Song, B. H., Blank, I. A., & Kao, J. C. (2026). Spurious alignment between large language models and brains can emerge from non-robust methods and overlooked confounds. Nature Communications, DOI: 10.1038/s41467-026-72253-7.

Harris, K. D. (2021). Nonsense correlations in neuroscience. bioRxiv, DOI: 10.1101/2020.11.29.402719.

Harris, K. D. & Thiele, A. (2011). Cortical state and attention. Nature Reviews Neuroscience, 12, 509–523.

Hastie, T., Tibshirani, R., & Friedman, J. (2009). The Elements of Statistical Learning. (Springer New York).

He, B., Yuan, H., Meng, J., & Gao, S. (2020). Brain–computer interfaces. In Neural Engineering, B. He, ed. (Springer International Publishing), pp. 131–183.

Horikawa, T. & Kamitani, Y. (2017). Generic decoding of seen and imagined objects using hierarchical visual features. Nature Communications, 8, 15037.

Horikawa, T., Tamaki, M., Miyawaki, Y., & Kamitani, Y. (2013). Neural decoding of visual imagery during sleep. Science, 340, 639–642.

Jun, J. J., Steinmetz, N. A., Siegle, J. H., Denman, D. J., Bauza, M., Barbarits, B., Lee, A. K., Anastassiou, C. A., Andrei, A., Aydın, C., Barbic, M., Blanche, T. J., Bonin, V., Couto, J., Dutta, B., Gratiy, S. L., Gutnisky, D. A., Häusser, M., Karsh, B., Ledochowitsch, P., Lopez, C. M., Mitelut, C., Musa, S., Okun, M., Pachitariu, M., Putzeys, J., Rich, P. D., Rossant, C., Sun, W.-l., Svoboda, K., Carandini, M., Harris, K. D., Koch, C., O’Keefe, J., & Harris, T. D. (2017). Fully integrated silicon probes for high-density recording of neural activity. Nature, 551, 232–236.

Kapoor, S. & Narayanan, A. (2023). Leakage and the reproducibility crisis in machine-learning-based science. Patterns, 4, 100804.

Karpowicz, B. M., Ali, Y. H., Wimalasena, L. N., Sedler, A. R., Keshtkaran, M. R., Bodkin, K., Ma, X., Rubin, D. B., Williams, Z. M., Cash, S. S., Hochberg, L. R., Miller, L. E., & Pandarinath, C. (2025). Stabilizing brain-computer interfaces through alignment of latent dynamics. Nature Communications, 16.

Kriegeskorte, N., Simmons, W. K., Bellgowan, P. S. F., & Baker, C. I. (2009). Circular analysis in systems neuroscience: the dangers of double dipping. Nature Neuroscience, 12, 535–540.

Kriegeskorte, N. & Wei, X.-X. (2021). Neural tuning and representational geometry. Nature Reviews Neuroscience, 22, 703–718.

Langdon, C., Genkin, M., & Engel, T. A. (2023). A unifying perspective on neural manifolds and circuits for cognition. Nature Reviews Neuroscience, 24, 363–377.

Lin, A., Witvliet, D., Hernandez-Nunez, L., Linderman, S. W., Samuel, A. D. T., & Venkatachalam, V. (2022). Imaging whole-brain activity to understand behaviour. Nature Reviews Physics, 4, 292–305.

Lotte, F., Bougrain, L., Cichocki, A., Clerc, M., Congedo, M., Rakotomamonjy, A., & Yger, F. (2018). A review of classification algorithms for eeg-based brain-computer interfaces: a 10 year update. Journal of Neural Engineering, 15, 031005.

Ma, X., Rizzoglio, F., Bodkin, K. L., Perreault, E., Miller, L. E., & Kennedy, A. (2023). Using adversarial networks to extend brain computer interface decoding accuracy over time. eLife, 12.

Mahalanobis, P. C. (1936). On the generalized distance in statistics. Proceedings of the National Institute of Sciences (Calcutta), 2, 49–55.

McLachlan, G. J. (1999). Mahalanobis distance. Resonance, 4, 20–26.

Mignacco, F., Krzakala, F., Lu, Y., Urbani, P., & Zdeborova, L. (2020). The role of regularization in classification of high-dimensional noisy Gaussian mixture. In Proceedings of the 37th International Conference on Machine Learning, H. D. III & A. Singh, eds., vol. 119 of Proceedings of Machine Learning Research, pp. 6874–6883. (PMLR).

Moscovich, A. & Rosset, S. (2022). On the cross-validation bias due to unsupervised preprocessing. Journal of the Royal Statistical Society Series B: Statistical Methodology, 84, 1474–1502.

Perich, M. G., Narain, D., & Gallego, J. A. (2025). A neural manifold view of the brain. Nature Neuroscience, 28, 1582–1597.

Petrucco, L. (2020). Zebrafish larvae lateral view. Zenodo. DOI: 10.5281/ZENODO.3926123.

Pillow, J. & Scott, J. (2012). Fully bayesian inference for neural models with negative-binomial spiking. In Advances in Neural Information Processing Systems, F. Pereira, C. Burges, L. Bottou, & K. Weinberger, eds., vol. 25. (Curran Associates, Inc.).

Pnevmatikakis, E. A., Soudry, D., Gao, Y., Machado, T. A., Merel, J., Pfau, D., Reardon, T., Mu, Y., Lacefield, C., Yang, W., Ahrens, M., Bruno, R., Jessell, T. M., Peterka, D. S., Yuste, R., & Paninski, L. (2016). Simultaneous denoising, deconvolution, and demixing of calcium imaging data. Neuron, 89, 285–299.

Poldrack, R. A., Huckins, G., & Varoquaux, G. (2020). Establishment of best practices for evidence for prediction: A review. JAMA Psychiatry, 77, 534.

Rosenblatt, M., Tejavibulya, L., Jiang, R., Noble, S., & Scheinost, D. (2024). Data leakage inflates prediction performance in connectome-based machine learning models. Nature Communications, 15, 1829.

Rumyantsev, O. I., Lecoq, J. A., Hernandez, O., Zhang, Y., Savall, J., Chrapkiewicz, R., Li, J., Zeng, H., Ganguli, S., & Schnitzer, M. J. (2020). Fundamental bounds on the fidelity of sensory cortical coding. Nature, 580, 100–105.

Saxena, S. & Cunningham, J. P. (2019). Towards the neural population doctrine. Current Opinion in Neurobiology, 55, 103–111.

Schneider, S., Lee, J. H., & Mathis, M. W. (2023). Learnable latent embeddings for joint behavioural and neural analysis. Nature, 617, 360–368.

Shim, M., Lee, S.-H., & Hwang, H.-J. (2021). Inflated prediction accuracy of neuropsychiatric biomarkers caused by data leakage in feature selection. Scientific Reports, 11, 7980.

Simon, R., Radmacher, M. D., Dobbin, K., & McShane, L. M. (2003). Pitfalls in the use of dna microarray data for diagnostic and prognostic classification. JNCI Journal of the National Cancer Institute, 95, 14–18.

Steinmetz, N. A., Aydin, C., Lebedeva, A., Okun, M., Pachitariu, M., Bauza, M., Beau, M., Bhagat, J., Böhm, C., Broux, M., Chen, S., Colonell, J., Gardner, R. J., Karsh, B., Kloosterman, F., Kostadinov, D., Mora-Lopez, C., O’Callaghan, J., Park, J., Putzeys, J., Sauerbrei, B., van Daal, R. J. J., Vollan, A. Z., Wang, S., Welkenhuysen, M., Ye, Z., Dudman, J. T., Dutta, B., Hantman, A. W., Harris, K. D., Lee, A. K., Moser, E. I., O’Keefe, J., Renart, A., Svoboda, K., Häusser, M., Haesler, S., Carandini, M., & Harris, T. D. (2021). Neuropixels 2.0: A miniaturized high-density probe for stable, long-term brain recordings. Science, 372, eabf4588.

Stone, M. (1974). Cross-validatory choice and assessment of statistical predictions. Journal of the Royal Statistical Society. Series B (Methodological), 36, 111–147.

Stringer, C., Michaelos, M., Tsyboulski, D., Lindo, S. E., & Pachitariu, M. (2021). High-precision coding in visual cortex. Cell, 184, 2767–2778.e15.

Tibshirani, R. (1996). Regression shrinkage and selection via the lasso. Journal of the Royal Statistical Society. Series B (Methodological), 58, 267–288.

Urai, A. E., Doiron, B., Leifer, A. M., & Churchland, A. K. (2022). Large-scale neural recordings call for new insights to link brain and behavior. Nature Neuroscience, 25, 11–19.

Verstynen, T. & Kording, K. P. (2023). Overfitting to ‘predict’ suicidal ideation. Nature Human Behaviour, 7, 680–681.

Vogelstein, J. T., Packer, A. M., Machado, T. A., Sippy, T., Babadi, B., Yuste, R., & Paninski, L. (2010). Fast nonnegative deconvolution for spike train inference from population calcium imaging. Journal of Neurophysiology, 104, 3691–3704.

Wang, E. Y., Fahey, P. G., Ding, Z., Papadopoulos, S., Ponder, K., Weis, M. A., Chang, A., Muhammad, T., Patel, S., Ding, Z., Tran, D., Fu, J., Schneider-Mizell, C. M., da Costa, N. M., Reid, R. C., Collman, F., da Costa, N. M., Franke, K., Ecker, A. S., Reimer, J., Pitkow, X., Sinz, F. H., & Tolias, A. S. (2025). Foundation model of neural activity predicts response to new stimulus types. Nature, 640, 470–477.

Wen, H., Shi, J., Zhang, Y., Lu, K.-H., Cao, J., & Liu, Z. (2017). Neural encoding and decoding with deep learning for dynamic natural vision. Cerebral Cortex, 28, 4136–4160.

Wolpaw, J. R. & McFarland, D. J. (2004). Control of a two-dimensional movement signal by a noninvasive brain-computer interface in humans. Proceedings of the National Academy of Sciences, 101, 17849–17854.

Ye, J., Collinger, J. L., Wehbe, L., & Gaunt, R. (2023). Neural data transformer 2: Multi-context pretraining for neural spiking activity. DOI: 10.1101/2023.09.18.558113.

Yu, B. M., Cunningham, J. P., Santhanam, G., Ryu, S. I., Shenoy, K. V., & Sahani, M. (2009). Gaussian-process factor analysis for low-dimensional single-trial analysis of neural population activity. Journal of Neurophysiology, 102, 614–635.

## References

Barabási, A.-L. & Albert, R. (1999). Emergence of scaling in random networks. Science, 286, 509–512.

Buitinck, L., Louppe, G., Blondel, M., Pedregosa, F., Mueller, A., Grisel, O., Niculae, V., Prettenhofer, P., Gramfort, A., Grobler, J., Layton, R., VanderPlas, J., Joly, A., Holt, B., & Varoquaux, G. (2013). API design for machine learning software: experiences from the scikit-learn project. In ECML PKDD Workshop: Languages for Data Mining and Machine Learning, pp. 108–122.

Combrisson, E. & Jerbi, K. (2015). Exceeding chance level by chance: The caveat of theoretical chance levels in brain signal classification and statistical assessment of decoding accuracy. Journal of Neuroscience Methods, 250, 126–136.

Fornito, A., Zalesky, A., & Bullmore, E. (2016). Chapter 10 - null models. In Fundamentals of Brain Network Analysis. (Academic press), pp. 355–381.

Ojala, M. & Garriga, G. C. (2010). Permutation tests for studying classifier performance. Journal of Machine Learning Research, 11, 1833–1863.

Váša, F. & Mišić, B. (2022). Null models in network neuroscience. Nature Reviews Neuroscience, 23, 493–504.

